# On the Mathematics of RNA Velocity I: Theoretical Analysis

**DOI:** 10.1101/2020.09.19.304584

**Authors:** Tiejun Li, Jifan Shi, Yichong Wu, Peijie Zhou

## Abstract

The RNA velocity provides a new avenue to study the stemness and lineage of cells in the development in scRNA-seq data analysis. Some promising extensions of it are proposed and the community is experiencing a fast developing period. However, in this stage, it is of prime importance to revisit the whole process of RNA velocity analysis from the mathematical point of view, which will help to understand the rationale and drawbacks of different proposals. The current paper is devoted to this purpose. We present a thorough mathematical study on the RNA velocity model from dynamics to downstream data analysis. We derived the analytical solution of the RNA velocity model from both deterministic and stochastic point of view. We presented the parameter inference framework based on the maximum likelihood estimate. We also derived the continuum limit of different downstream analysis methods, which provides insights on the construction of transition probability matrix, root and ending-cells identification, and the development routes finding. The overall analysis aims at providing a mathematical basis for more advanced design and development of RNA velocity type methods in the future.

## 1 Introduction

Single-cell RNA sequencing (scRNA-seq) is a rapid maturing technique, which makes the elaborate study of biological processes in the single cell resolution possible [48, 56]. The rich and diverse scRNA-seq datasets are revealing to us the mysteries of stem cell differentiation [52], heterogeneity in multicellular organisms [23], cancer cell dissection [7, 35, 62], drug discovery [21, 57], etc. Every year, a swarm of analysis tools are produced by researchers all over the world [40, 60]. Some popular choices include the clustering tools [6, 26], trajectory inference tools [20, 38, 40, 45, 49, 51], and energy landscape tools [24, 42, 44, 61], etc.

The characterization of stemness and lineage of the cells is a fundamental question in developmental biology. Although some practical indices, such as the signalling entropy and Markov chain entropy [45, 49], etc., are proposed to quantify the stemness of different cells in the scRNAseq data analysis, they are more or less heuristic in nature. Recently, another promising method, the RNA velocity [27], was proposed to address this issue based upon the fact that the nascent (unspliced) and mature (spliced) mRNA can be distinguished in common singlecell RNA-seq protocols, such as SMART-seq2 [36], Dropseq [30] and 10X genomics [64]. Thus, the relative abundance of unspliced and spliced mRNA are utilized to infer a velocity of each cell in the spliced mRNA abundance space, and predict the tendency of transition from one cell to another according to the RNA velocity model [27]. Improved methods in kinetic modeling, parameter inference and downstream analysis have been subsequently proposed [3, 39], showing the potential of RNA velocity to quantify the stemness of cells in a rational way.

Despite the fruitful results and promising applications of RNA velocity, it is of prime importance to understand the rationale underlying the algorithm design, as well as the subtle differences between different proposals from the mathematical point of view. For instance, when constructing the cell-cell stochastic transition probability matrix from RNA velocity, La Mano et al. [27] and Qiu et al.[39] used the correlation scheme in the velocity kernel, while the cosine scheme was proposed in [3]. In the recent version of dynamo package [37], a scheme with local kernels [4] of diffusion was also utilized. In spite of their intuitive plausibility, the theoretical implications of different kernels demands further investigation. In addition, a tracking strategy of root and ending cells has been applied based on forward and backward diffusions [3, 27], whose theoretical basis remains to be established. The resolve of these puzzles based on a formal mathematical study will not only shed light on these theoretical problems, but also lead to a deeper comprehension of the RNA velocity and inspire further rational design of more delicate RNA velocity models. The current paper is devoted to this purpose.

In this work, we will present a thorough mathematical study on the whole process of RNA velocity model from kinetic model derivation, parameter inference algorithm to the downstream dynamical analysis. Our analysis will contribute insights toward several fundamental questions regarding RNA velocity and relevant downstream analysis, including:

- How to derive the deterministic and stochastic kinetic models of RNA velocity, and find analytical solutions?
- How to build the maximum likelihood estimator (MLE) of the parameters, built on the exact solution of stochastic RNA velocity model?
- How can the discrete cellular transition dynamics inferred from RNA velocity be rigorously associated the continuous dynamical system model in cell-fate decision?
- What is the essential difference between the choice of correlation, cosine or innerproduct scheme in the velocity kernel for the cellular transition matrix?
- What is the implication to replace the Gaussian scheme with k-nearest neighbor (kNN) scheme in the diffusion kernel?
- Why is the backward and forward diffusion strategy effective in detecting root and ending cells of development?
- How to rationally construct developmental trajectories based on RNA velocity with mathematical theory, beyond illustrating arrows and streamlines in the reduced dimension space?

We will focus on the formal mathematical analysis in the current paper and leave detailed computational comparisons and improvements in the continued publication [28]. To the best knowledge of the authors, this is the first attempt on studying the mathematics of RNA velocity in a complete manner. We hope it will provide a mathematical basis for further development of RNA velocity type methods in the future.

The rest of the paper is organized as follows. In Section 2, we show the mathematical derivations of both deterministic and stochastic kinetic models of RNA velocity, and derive the associated analytical solutions. In Section 3 we will revisit the existing algorithms to infer parameters in RNA velocity models, and present a novel maximum likelihood estimation of parameters originated from the exact solution of stochastic models. In Section 4 we focus on the dynamical system analysis based on RNA velocity, deriving the continuum limit of discrete transition probabilities with various kernels, demonstrating the mathematical rationale for the existing strategy of root/ending cells detection, and providing a new method to construct development trajectories with RNA velocity through the well-established transition path theory. Finally we give the conclusion and discussions in Section 5. Some analysis details, such as the almost sure type convergence order of the kNN radius, are left in the Appendix.

## 2 Models of RNA Velocity

The key point of the RNA velocity model of single cells is that one can identify the abundance of the precursor (unspliced) and mature (spliced) mRNA from the single-cell RNA-seq data, which provides the information on the time-dependent evolution rate of the mRNA abundance by incorporating appropriate dynamical models.

Denote by *u* and *s* the abundance of the unspliced and spliced mRNA, respectively. In its simplest form, the transcriptional dynamics of the mRNA velocity can be described as the reaction pathways shown in Table 1 [27]. We assume the production of *u* is dictated by a transcriptional induction or repression with parameter *α*^on^ or *α*^off^, respectively. The unspliced mRNA, *u*, is then transformed into the spliced form with rate *β*, and the spliced mRNA is eventually degradated with rate *γ*. We remark that the above statement must be understood for single gene, i.e. the parameters (*α*^on/off^, *β, γ*) should be replaced by 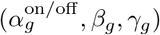 when we consider the dynamics for a specific gene *g*. But we will omit the *g*-dependence of the parameters for brevity if not necessary. In the current stage, we assume that there are no interactions among different genes.

**Table 1:**
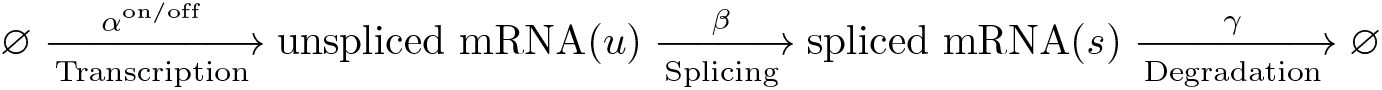
Schematics of the dynamics for the RNA velocity model.

The task in this section is to study the explicit solution and related analytical properties of the forward mRNA velocity model in both deterministic and stochastic forms, given the dynamical parameters (*α*^on/off^, *β, γ*).

### 2.1 Deterministic Model

The deterministic model of the reaction dynamics shown in Table 1 has the form (2.1)-(2.2) by the law of mass action:

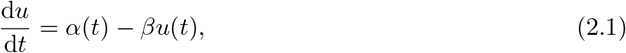

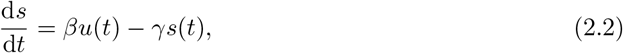

where *t* ≥ 0, (*u*(*t*), *s*(*t*))|_*t*=0_ = (*u*_0_, *s*_0_), and

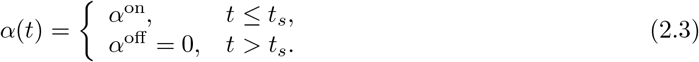

Here *t*_*s*_ is the switch time of the transcriptional process.

The term defined through Eq. (2.2):

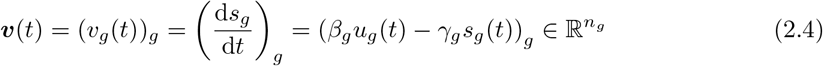

is the RNA velocity of each cell, where *g* = 1 : *n*_*g*_ in Eq. (2.4) and *n*_*g*_ is the number of considered genes in the RNA-seq data. Note that the velocity ***v*** only depends on the state (*u, s*), but not the absolute magnitude of *t*, given the rate parameters, since the considered system is autonomous.

#### Explicit solution

We will study the results for the cases *β* ≠ *γ* and *β* = *γ*, respectively.

*Case 1: β* ≠ *γ.* The analytical solution to (2.1)-(2.2) in the *on* stage with rate *α*^on^ = *α* is

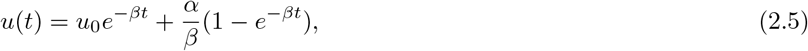

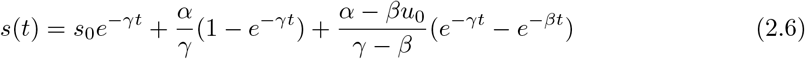

for *t* ≤ *t*_*s*_. Usually, we suppose (*u*_0_, *s*_0_) = (0, 0), then we have

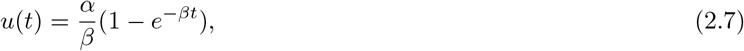

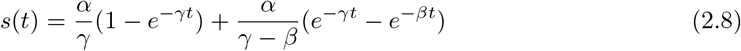

for *t* ≤ *t*_*s*_. Define the switch state by

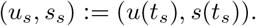

Then in the *off* stage, we have the solution

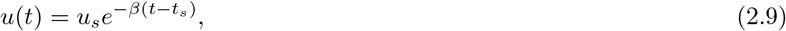

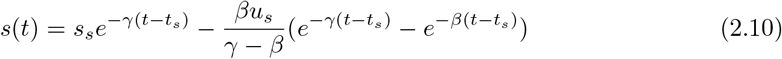

for *t > t*_*s*_. It is straightforward that *u*(*t*), *s*(*t*) *>* 0 for any finite *t >* 0.

*Case 2: β* = *γ.* The analytical solution to (2.1)-(2.2) in the *on* stage is

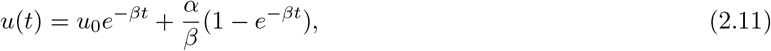

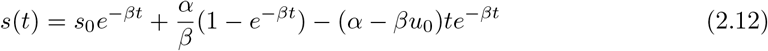

for *t* ≤ *t*_*s*_. When (*u*_0_, *s*_0_) = (0, 0), we have

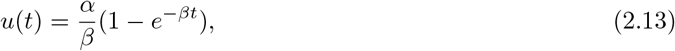

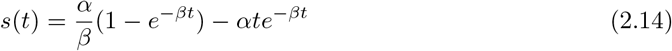

for *t* ≤ *t*_*s*_. And in the off stage

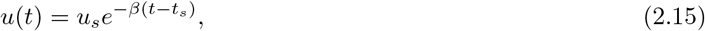

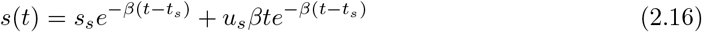

for *t > t*_*s*_. It is straightforward to note that the solution (2.11)-(2.12) is indeed the limit of the solution (2.5)-(2.6) as *γ* → *β*.

#### Steady State

The steady state in the on stage is (*u*\*, *s*\*) = *α/*(*β, γ*), and the steady state in the off stage is simply (*u*_*_, *s*_*_) = (0, 0).

#### Scale Invariance

It is important to note that the system (2.1)-(2.2) has the following scale invariance property, i.e. if we define the parameter *θ* = (*θ*_*r*_, *t*_*s*_), where the rates *θ*_*r*_ = (*α, β, γ*), then the solution satisfies

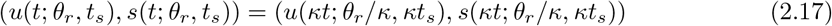

for any scaling parameter *κ >* 0. This scale invariance indicates the degeneracy of the inference problem. That is, to ensure the well-posedness of the inference on parameter *θ*, we should fix the time scale of the system. For example, one can consider the dynamics (2.1)-(2.2) within a fixed period [0, *T*], where *t*_max_ = *T*. We remark that the choice of the degree of freedom does affect the magnitude of RNA velocity (2.4) up to a multiplicative constant.

### 2.2 Stochastic Model

In the stochastic model, the system state (*u*(*t*), *s*(*t*)) ∈ ℕ^2^ is a stochastic process and we are interested in the evolution of its probability mass function denoted by

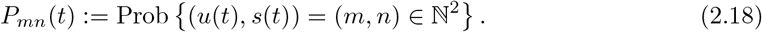

The stochastic model of the reaction dynamics shown in Table 1 is given by the following chemical master equation (CME) [18]

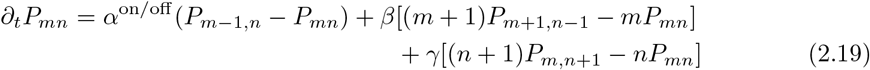

with initial condition 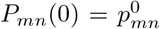. In the on or off stage, the production rate of *u*, *α*^on/off^, will be set as *α*^on^ = *α* or *α*^off^ = 0, respectively. We will study the analytical solution of (2.19) in different cases.

#### Scale Invariance

Similar to the deterministic case, the solution *P*_*mn*_(*t*) has the scale invariance property

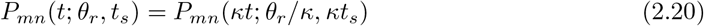

for any scaling parameter *κ >* 0.

##### 2.2.1 On Stage with Zero Initial Value

In the on stage, i.e. *t* ≤ *t*_*s*_, we have the CME

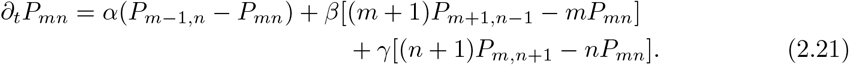

We will first study the case (*u*(0), *s*(0)) = (0, 0), i.e. with initial distribution *P*_*mn*_(0) = *δ*_*m*0_*δ*_*n*0_, where

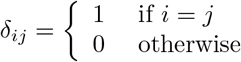

is the Kronecker’s delta-function. The general cases are left in Section 2.2.3. Since the rate functions are all linear in *m* and *n*, we will employ the idea of moment generating function to solve (2.21) [43].

###### Theorem 2.1

(Analytical Distribution in the On Stage). *With initial distribution P*_*mn*_(0) = *δ*_*m*0_*δ*_*n*0_, *the solution of* Eq. (2.21) *is*

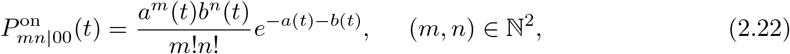

*where*

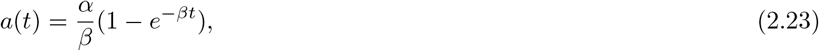

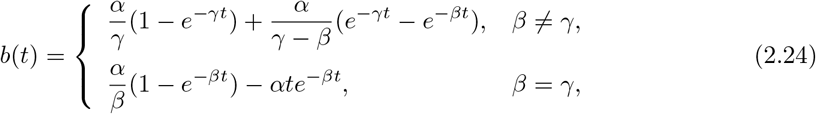

*and the notation 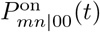 stands for the transition probability from state* (0, 0) *to* (*m, n*).

*Proof.* Consider the moment generating function *F*(*y, z, t*) = Σ_*m,n*_ *y*^*m*^*z*^*n*^*P*_*m,n*_(*t*). Then we have

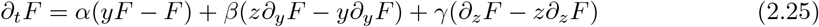

with initial condition *F*_0_(*y, z*) = Σ_*m,n*_*y*^*m*^*z*^*n*^*P*_*mn*_(0). Under the zero initial value condition on (*u*(*t*), *s*(*t*)), we have *F*_0_ = 1.

Introduce the change of variable

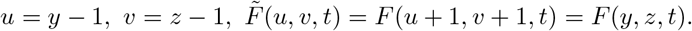

Then from (2.25) we get

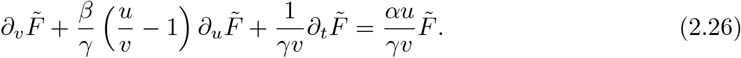

By the method of characteristics, we introduce the auxiliary variable *r*:

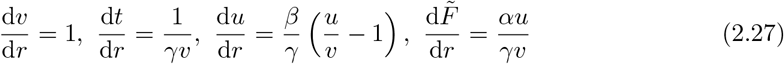

with initial condition 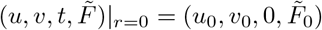, where 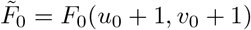.

*Case 1: β* ≠ *γ*. Solving (2.27), we get *v* = *r* + *v*_0_, d*v* = d*r, t* = *γ*^−1^ ln *v/v*_0_, and correspondingly

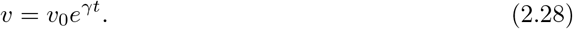

For *u*, we obtain

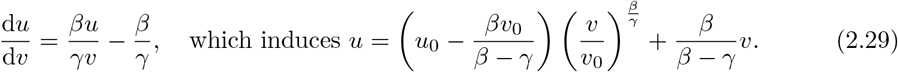

Define *a*_0_ ≔ *u*_0_ − (*β* − *γ*)^−1^*βv*_0_, we get

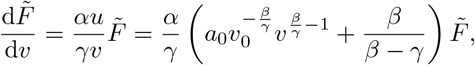

thus

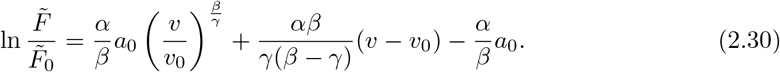

With initial distribution *P*_*m,n*_(0) = *δ*_*m*0_*δ*_*n*0_, we have 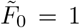. Combining Eqs. (2.28), (2.29), (2.30) and the definition of *a*_0_, we get

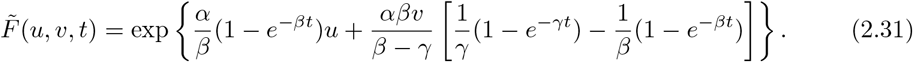

After suitable manipulation, we obtain

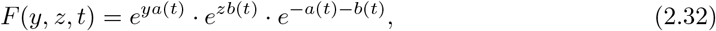

where *a*(*t*) and *b*(*t*) are defined in (2.23) and (2.24) when *β* ≠ *γ*, respectively.

*Case 2: β* = *γ.* We can show that (2.29) will be replaced by

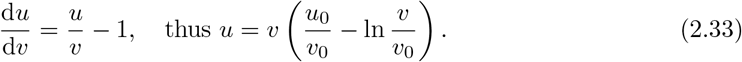

After suitable derivations, we get

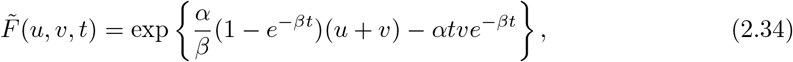

and finally we have the same formula (2.32), while *a*(*t*) and *b*(*t*) are defined in (2.23) and (2.24) when *β* = *γ*, respectively.

Therefore,

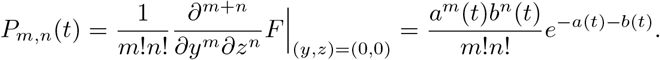

In summary, *u*(*t*) and *s*(*t*) are independently Poisson distributed with mean *a*(*t*) and *b*(*t*), respectively.

###### Remark 2.1.

*It is not surprising to observe that the mean a*(*t*), *b*(*t*) *are exactly the abundance of u and s in Eqs.* (2.7)-(2.8) *or* (2.13)-(2.14) *in the deterministic model, which is well-known due to the linearity of the rates. However, Theorem 2.1 further states that u and s are independently Poisson distributed, which is not a straightforward result.*

#### Invariant distribution

It is obvious that the invariant distribution of (*u, s*) in this case is independent Poisson with parameters (*u*\*, *s*\*).

##### 2.2.2 Off Stage with General Initial Data

We will study the off stage case in this section. Now we have *α*^off^ = 0 and the CME is

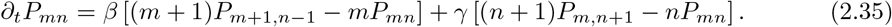

We first consider the case with initial value (*u*(0), *s*(0)) = (*M, N*).

###### Theorem 2.2

(Analytical Distribution in the Off Stage). *Define*

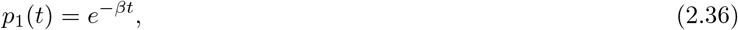

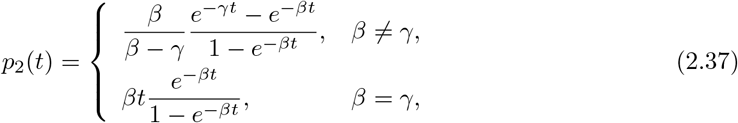

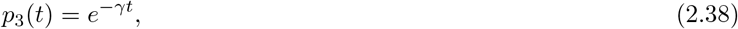

*and q*_*i*_(*t*) = 1 − *p*_*i*_(*t*) *correspondingly. We have p*_*i*_(*t*), *q*_*i*_(*t*) ∈ [0, 1] *for i* = 1, 2, 3. *Then, with initial distribution P*_*mn*_(0) = *δ*_*mM*_*δ*_*nN*_, *the solution of* Eq. (2.35) *has the form*

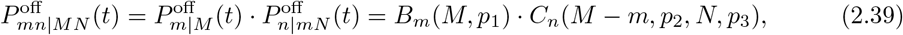

*where m* ≤ *M, n* ≤ *N* + *M* − *m*,

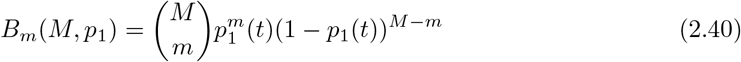

*is the probability of the binomial distribution* 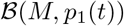, *and*

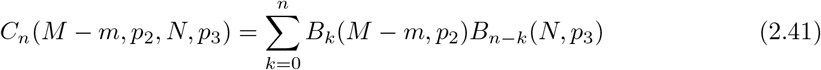

*is the probability of the sum of two independent binomials* 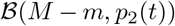 *and* 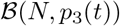. *We take the convention that B*_*k*_(*M* − *m, p*_2_) = 0 *if k > M* − *m.*

*Proof.* First let us show that *p*_2_(*t*) ∈ [0, 1]. When *β* ≠ *γ*, we have

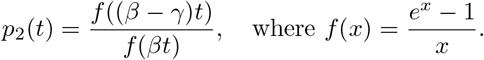

We have *f′*(*x*) = (*e*^*x*^(*x* − 1) + 1)*/x*^2^ ≥ 0 since *g*(0) = 0, and *g′* (*x*) *>* 0 for *x >* 0 and *g′* (*x*) *<* 0 for *x <* 0, where *g*(*x*) ≔ *e*^*x*^(*x* − 1) + 1. The case *β* = *γ* is trivial by observing that the function *x/*(*e*^*x*^ − 1) ∈ [0, 1] for *x* ≥ 0.

Next we derive the distribution *P*_*mn*_. Similar to the proof of Theorem 2.1, for moment generating function *F*(*y, z, t*) = Σ_*m,n*_ *y*^*m*^*z*^*n*^*P*_*m,n*_(*t*), we have

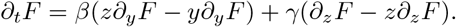

Similarly define

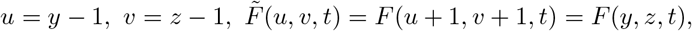

we obtain

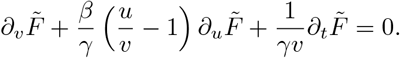

Introduce the parameter *r*, we get by the method of characteristics

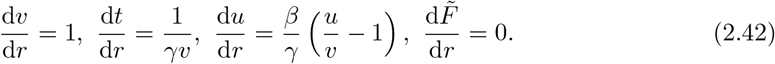

*Case 1: β* ≠ *γ.* Similar derivation shows

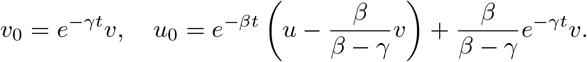

We obtain

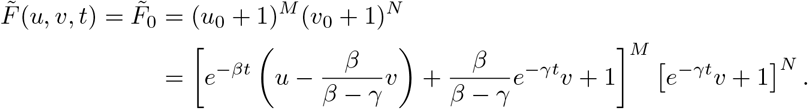

After suitable manipulations, we get

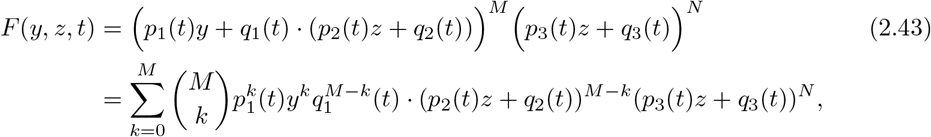

which exactly has the probabilistic interpretation as shown in Eqs. (2.40)-(2.41).

*Case 2: β* = *γ.* We can show that

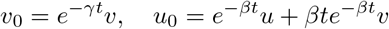

in this case. Substitute into 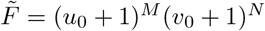, we get the same equation (2.43) but with the *p*_*i*_(*t*) in the *β* = *γ* case.

###### Remark 2.2.

*In the off stage, the unspliced mRNA u obeys the binomial distribution. Given u*(*t*) = *m, the conditional distribution of the spliced mRNA s is characterized by the sum of two independent binomial random variables. Intuitively, s is comprised of two parts: the new spliced mRNA counts generated from u and the non-degradated spliced mRNA counts from the initial state. It is also natural to observe that the mean of* (*u*(*t*), *s*(*t*)) *is* (*Mp*_*1*_(*t*), *Mq*_*1*_(*t*)*p*_*2*_(*t*)+*Np*_*3*_(*t*)), *which is essentially* (2.9)-(2.10).

###### Corollary 2.1.

*When the initial distribution* 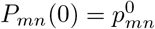, *the solution of* Eq. (2.35) *is*

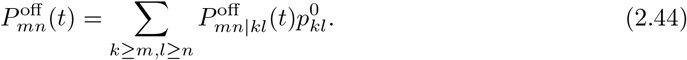

*for* (*m, n*) ∈ ℕ^2^.

#### Invariant distribution

It is straightforward that the the invariant distribution in the off stage is simply *P*_*mn*_(∞) = *δ*_*;m*0_*δ*_*n*0_ as *t* → ∞.

##### 2.2.3 On Stage with General Initial Data

###### Corollary 2.2.

*When the initial distribution* 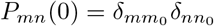, *the solution of* Eq. (2.21) *is*

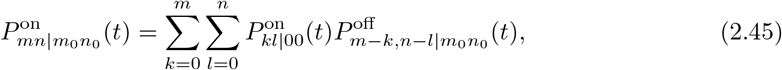

*which is the convolution of the distributions* 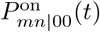 *and* 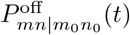. *We adopt the convention that* 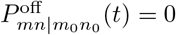. *if m >m*_0_ *or n > n*_0_

*Proof.* To check the result, we only need to note that 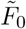 in (2.30) will be replaced by 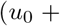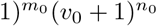. Thus we have

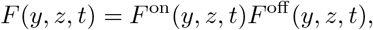

where *F*^on^(*y, z, t*) and *F*^off^(*y, z, t*) are defined as in (2.32) and (2.43), respectively. This naturally yields to the transition probability (2.45).

###### Remark 2.3.

*The above result leads to a recipe to directly generate samples from* 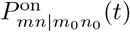:

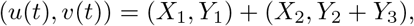

*where X*_1_, *Y*_1_, *X*_2_, *Y*_3_ *are independent random variables with* 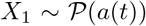, 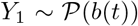, 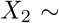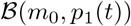, 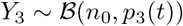, *and* 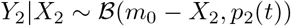.

###### Corollary 2.3.

*When the initial distribution* 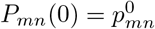, *the solution of* Eq. (2.21) *is*

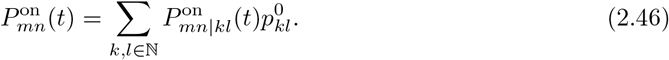

*for* (*m, n*) ∈ ℕ^2^.

## 3 Inference of RNA velocity

In this section, we will study the inverse problem: the inference of the parameters in the RNA velocity model from the data. We will mainly revisit the proposals pursued in [3, 27] and briefly mention our new approach [58], which utilizes the full stochastic model to do the inference.

### 3.1 Steady State Model

The steady state model was first considered in [27]. In this model, one assumes that the on stage lasts sufficiently long so the state of the system is close to the steady state of the dynamical system (2.1)-(2.2). Therefore, the upper-right corner points in the (*u, s*)-plot can be approximated as steady states. In steady states, we have

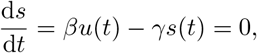

which means that the mRNA synthesis and degradation are in balance. This balance condition in the steady state can be utilized to approximate the ratio of degradation and splicing rates via least squares fitting as

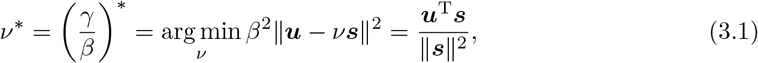

where ***u, s*** are vectors with components corresponding to the cells in the upper-right corner points in the (*u, s*)-plot for each gene. If further assuming that *β* = 1 across all genes in [27] via scale invariance argument, the RNA velocity is then estimated as

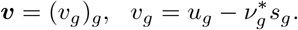

Though original, simple and successful, the above steady state model and the treatment with *β* = 1 for all genes are not good enough assumptions in many cases. In fact, setting the splicing rates *β* = 1 for all genes is actually wrong according to the scale invariance property of the system, which only permits one degree of freedom to be adjusted. These drawbacks call for more robust and accurate estimation methods for the RNA velocity.

### 3.2 EM Algorithm for the Transient Models

In this subsection, we will revisit and study the parameter inference using EM algorithm for transient models. The most related references on this aspect are [3, 58]. To ensure the computational feasibility, we also employ suitable approximations, which will be stated in the corresponding places below.

#### 3.2.1 Basic Framework of EM

Given the observed data 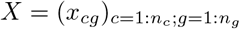, where *x*_*cg*_ = (*u*_*cg*_, *s*_*cg*_) for cell *c* and gene *g*, we want to maximize the log-likelihood

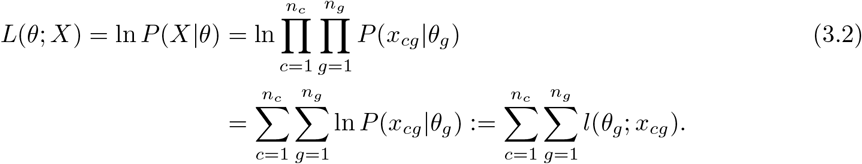

Note that the abundance *x*_*cg*_ depends on time *t*_*cg*_ and the switch state *x*_*s,cg*_ = (*u*_*s,cg*_, *s*_*s,cg*_), which are not observables, we indeed encounter a hidden variable problem. It is natural to utilize the EM algorithm to do the inference.

Let us introduce the latent variable *h*_*cg*_ = (*t*_*cg*_, *x*_*s,cg*_) for *c* = 1 : *n*_*c*_, *g* = 1 : *n*_*g*_, where *t* is the latent time and *x*_*s*_ is the latent switch state. Then the log-likelihood function can be written as

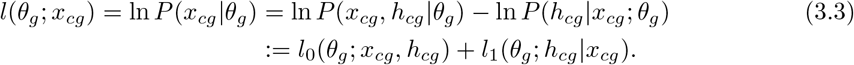

Similarly denote the sum of *l*_0_(*θ*_*g*_; *x*_*cg*_, *h*_*cg*_) and *l*_1_(*θ*_*g*_; *h*_*cg*_|*x*_*cg*_) with respect to *c, g* as *L*_0_(*θ*; *X, h*) and *L*_1_(*θ*; *h*|*X*), respectively. Then

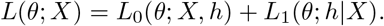

Taking conditional expectation with respect to the distribution of *h*|*X* given parameter *θ′*, we get

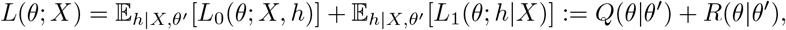

where

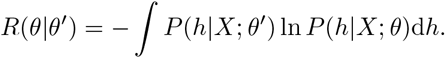

The above formulation is the basis of the well-known EM algorithm [10], which can be stated as below.

##### Algorithm 1 EM Algorithm for the RNA Velocity Model.

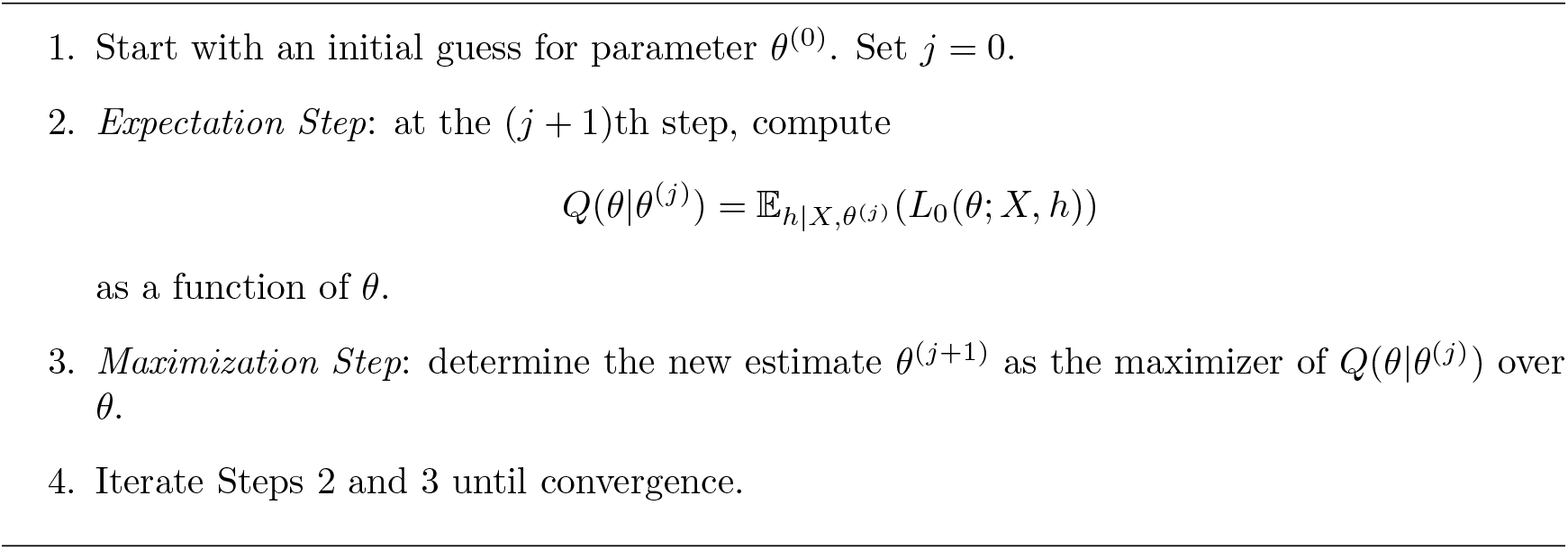

It is a classical result that the EM iterations never decreases the log-likelihood *L*(*θ*; *X*). In fact, if *θ* maximizes *Q*(*θ*|*θ′*), we have

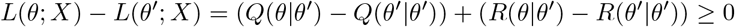

since *R*(*θ|θ′*) *R*(*θ′|θ′*) = *D*_KL_(*P*(*h*|*X*; *θ′*)||*P*(*h*|*X*; *θ*))≥ 0 by the non-negativity of Kullback-Leibler divergence [9]. This feature guarantees the local convergence of EM iterations.

#### 3.2.2 EM for the Deterministic Model

For the deterministic RNA velocity model, the latent variable *h* can be reduced to *t* since the switch state *x*_***s***_ = *x*(*t*_***s***_; *θ*) is uniquely determined. So we will replace *h* with *t* in (3.3) in the deterministic setup. If we assume the observation noise is Gaussian with mean 0 and variance *σ*^2^ for all cells and genes, and the sampling time *t*_*cg*_ is uniformly distributed in a fixed period [0, *T*], we have

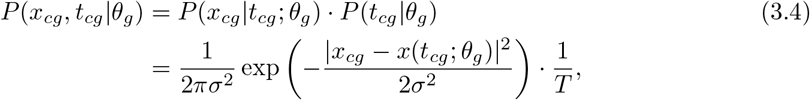

where *x*(*t*_*cg*_; *θ*_*g*_) is the solution of (2.1)-(2.2) at time *t*_*cg*_ with parameter *θ*_*g*_. Then

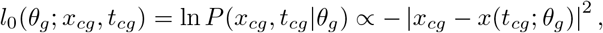

and

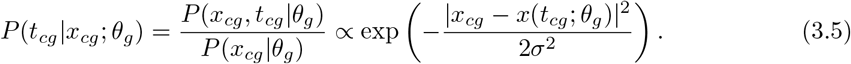

So we obtain

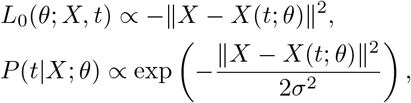

where 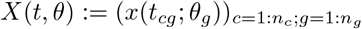.

According to EM Algorithm, we have

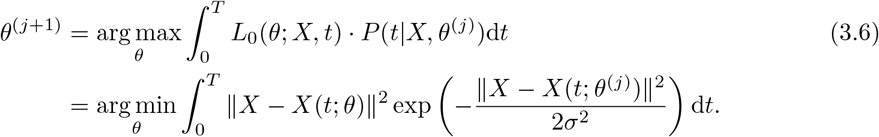

In the small noise limit regime, i.e. *σ* → 0, by Laplace asymptotics [2], we get

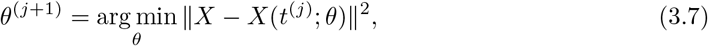

while

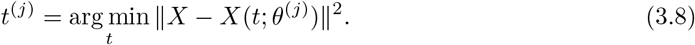

It forms an iteration between the parameter *θ* and the latent time *t*. Below we discuss more detailed procedure in (3.7)-(3.8).

##### Update of *t*

Two different models can be utilized in the update step (3.8), which we term the *independent-t model* and *uniform-t model* below. The two different choices lead to different computational complexity. We further assume that the switch time 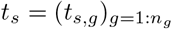 is only gene dependent throughout the transient model estimations.

###### Independent-t model

In this model, we permit the time *t*_*cg*_ for different *g* to be different, i.e. for a specific cell *c*, 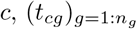 are independent. So we can estimate *t*_*cg*_ for each *c* and *g* separately.

The fact, that the estimation of 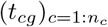 for different *g* can be separated, tells that we only need to consider a fixed *g*. Given *θ*_*g*_ and 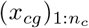, to estimate the optimal 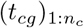, we classify the state of cell *c* into the on state if *t*_*cg*_ ≤ *t*_*s,g*_ or off state otherwise. Define the objective function

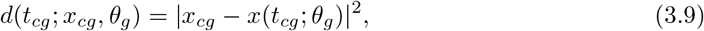

which has only two piecewise smooth parts determined by *t*_*s,g*_. We first compute the optimal 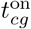 and 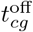 by assuming the cell *c* is in on or off stage, respectively with

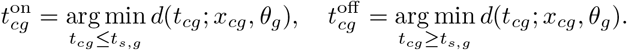

This optimization is feasible since we know the analytical form of *x*(*t*_*cg*_; *θ*_*g*_). The optimal estimation of *t*_*cg*_ is then obtained by

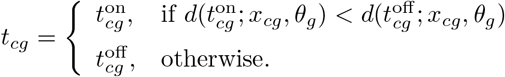

The computational complexity in this setup is 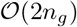 in terms of the analytical function evaluations of *x*(*t*; *θ*).

###### Uniform-t model

In this model, we require that the time *t*_*cg*_ for different genes are consistent, i.e. *t*_*cg*_ = *t*_*c*_ for *g* = 1 : *n*_*g*_. This setup is more reasonable in reality, however, it brings difficulty into the optimization.

In the uniform-*t* model, the objective functions (3.9) are no longer separated for different genes. Instead, we should consider the minimization of

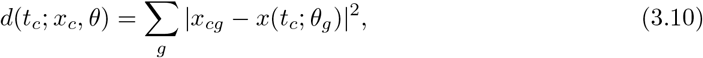

which generally has *n*_*g*_ + 1 piecewise smooth parts determined by 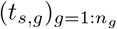. We can first sort the switch time *t*_*s*_ like

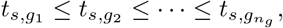

and next compute 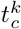 by miminizing (3.10) in each subinterval 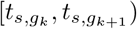 for *k* = 0, 1, *..., n*_*g*_ with 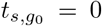 and 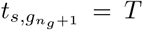. Finally we can obtain an optimal *t*_*c*_ which minimizes the objective functions 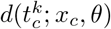.

The computational complexity in this setup is 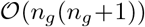 compared with the independent-*t* model.

##### Update of *θ*

For different genes *g*_1_ and *g*_2_, the update of 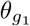 and 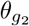 are independent according to (3.7). So we fix *g* and consider *θ*_*g*_ = (*α*_*g*_, *β*_*g*_, *γ*_*g*_, *t*_*s,g*_). There is no difference on the update of *θ*_*g*_ with independent-*t* model or uniform-*t* model because of the independence. We will only use the independent-*t* model formulation for illustration in the following text.

Define the objective function

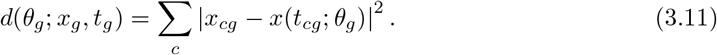

The minimization of (3.11) can be performed in different ways. One possible approach is to do the optimization with respect to the rates (*α*_*g*_, *β*_*g*_, *γ*_*g*_) and the switch time *t*_*s,g*_ alternatively, due to the non-smoothness induced by *t*_*s,g*_. Another approach is to reduce (3.11) into a new function of rates only

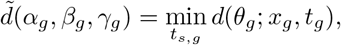

and then optimize 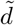 directly. In any case, the objective function is a sum of *n*_*c*_ terms with non-smoothness induced by *t*_*s,g*_. The optimization is not easy and usually stuck in the local minimum with local convergence methods.

###### Remark 3.1.

*We remark here that the choice of P*(*t*_*cg*_|*θ*_*g*_) = 1*/T can be altered, e.g. the uniform distribution along the curve* (*u*(*t*), *s*(*t*))*t*∈[0*,T*] *or other proposals, However, this will increase the computational complexity. When other choices of the observation noise are taken, the relative scales of the noise on different genes/cells still remain in the norm ||·|| in* (3.7)-(3.8) *even if we consider the zero noise limit regime.*

The above EM algorithm with independent-*t* model in the zero noise limit regime is utilized in [3].

#### 3.2.3 EM for the Stochastic Model

It is natural to consider the inference of the stochastic RNA velocity model. In the stochastic setup, the randomness of the switch state *x*_*s*_ should be incorporated into the full likelihood. Similarly assume that the observation noise is Gaussian with mean 0 and variance *σ*^2^ for all cells and genes, and the uniform distribution on the sampling time *t*_*cg*_ in a fixed period [0, *T*]. We have

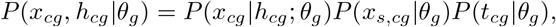

where

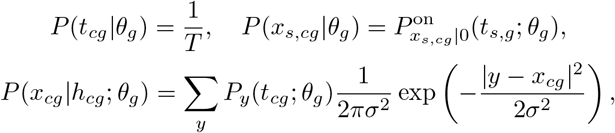

and

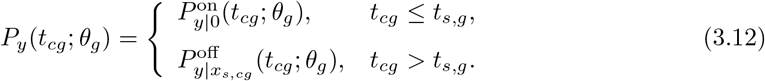

In the zero noise limit *σ* → 0, we get

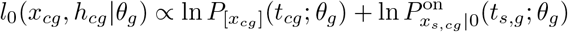

in the leading order, where [*x*] is the Gaussian nearest integer function. So we have

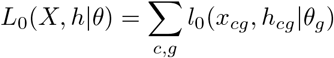

and correspondingly

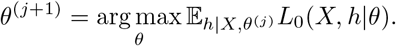

The optimization of the above formulation is not trivial. We leave further algorithmic constructions and practical applications to our continued publication [58].

## 4 Dynamical Analysis Based on RNA Velocity

After estimating the rates *θ*_*r*_ = (*α*_*g*_, *β*_*g*_, *γ*_*g*_)_*g*_ for each gene, one can then compute the RNA velocity 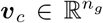 for each cell *c* = 1 : *n*_*c*_ via (2.4). One key component of the downstream analysis is to identify the source (stem cell) and sinks (differentiated cells) in the development process based on the obtained RNA velocities. We will discuss the related mathematics behind different proposals and give the rationale for this step. The continuum limit for various velocity kernels and the route-finding algorithm based on transition path theory will also be discussed.

### 4.1 Dynamical System View of Cell-Fate Development

Given a single cell, let 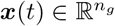 denote its gene expression profile (more generally, its *state* in cell-fate development) at time *t*. The evolution of ***x***(*t*) can be described by a dynamical system model using the ordinary differential equation (ODE)

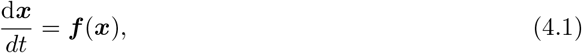

where the vector field 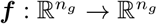 is determined by the gene regulatory kinetics.

Since the gene expression process is subject to both extrinsic and intrinsic noise [47], it is also common to model the cell-fate transition dynamics with stochastic ordinary differential equations (SDE)

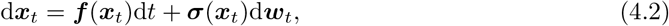

where ***w***_*t*_ ∈ ℝ^*k*^ denotes the standard Wiener process, representing the noise from *k* reaction channels or fluctuating sources, and the matrix 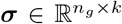 corresponds to the noise strength. The model is well-known as the chemical Langevin equation [19] with the appropriate coupling of ***f*** and *σ*.

As noted in [39], the RNA velocity for single cells can be incorporated in such dynamical system viewpoint to study the underlying cellfate dynamics. Currently, there are two lines of approaches to define the dynamical system utilizing RNA velocity in existing literatures:

1. The *continuous* dynamics approach [39]. In this approach, one fits a vector field 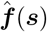 defined on the continuous space, such that 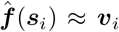 for each single cell, where ***v***_*i*_ are the estimated RNA velocities. Based on the inferred vector field 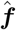, one can investigate the long-term dynamical properties of (4.1) or (4.2) to model the cell-fate development. The relevant important concepts include:

- Meta-stable states of cell fates development [22], corresponding to the attractors ***x***\* of (4.1) such that 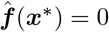.
- Energy landscape of cell fates development [65], which is the realization of Waddington’s metaphor [53]. Given the stochastic dynamics (4.2), among the different proposals for constructing energy landscape, one popular choice is the potential landscape [54] defined as follows. Note that evolution of probability density *p*(***x**, t*) in (4.2) can be described by the Fokker-Planck equation 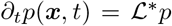, where 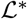 denotes the conjugate operator of the infinitesimal generator 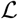 of SDE (4.2) such that

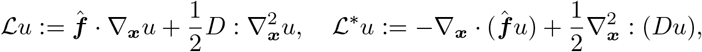

where *D*(***x***) = ***σσ***^T^ and *A* : *B* denotes Σ_*i,j*_ *A*_ij_*B*_*ij*_. The steady-state probability distribution *p*^*ss*^(***x***) satisfies 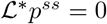, and the potential landscape of the system can be defined as *ϕ*(***x***) = − ln *p*^*ss*^(***x***).
2. The *discrete* dynamics approach [3, 27]. With this popular proposal, one utilizes the RNA velocity *v*_*i*_ to construct a Markov Chain defined on individual cells, with transition probabilities *P*(*s*_*i*_, *s*_*j*_) satisfying 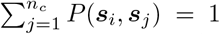. Ideally, the constructed Markov Chain should approximate the continuous dynamics (4.1) or (4.2) in discrete sense. Typical applications of such discrete dynamics include calculating the steady-state distribution of the system, and detecting roots and ending cells during development.

A fundamental theoretical question regarding the two approaches, which is also of noticeable mathematical interest, lies in the *consistency* issue between the two proposals. To be more specific, could the constructed discrete dynamics (with transition probabilities defined on individual data points) *converge* into the correct continuous dynamical systems, under appropriate limit regimes? In the next subsection, we will conduct a rigorous analysis on the different continuum limits of Markov chain dynamics, which are constructed with various choices of velocity kernels in defining transition probabilities.

### 4.2 Defining Transition Probabilities Among Cells

Let 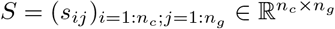 be the gene expression matrix of the spliced mRNA. We also denote

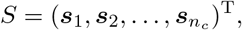

where 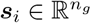 for *i* = 1 : *n*_*c*_. For ease of notation, we also use *d* = *n*_*g*_ for short in the following analysis.

To define the transition probability among different cells, one should take into account the randomness introduced by the unknown facts and the directed transition associated with the RNA velocity [3, 27]. The transition between two cells usually involves drift and diffusion. For the diffusion part, we consider the following two different *diffusion kernels*:

- Gaussian kernel

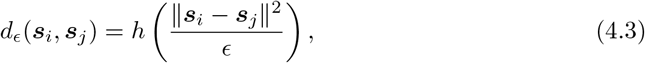
- kNN (k-Nearest Neighbor) kernel

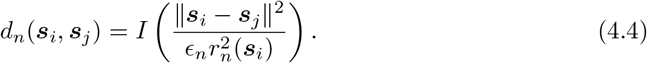

Here *h*(·) in (4.3) is a function with exponential decay, say *h*(*x*) ~ exp(−*x*) as *x* → ∞, *I*(·) in (4.4) is an indicator function with *I*(*r*) = 1 for |*r*| ≤ 1 and 0 otherwise, and 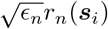 is the location-dependent distance to the *k*_*n*_th nearest neighbor of ***s***_*i*_ given *n* sample points (*n* = *n*_*c*_ in the above setup).

The kNN kernel can be reformulated as similar form in (4.3). Following [50], we have the kNN density estimate

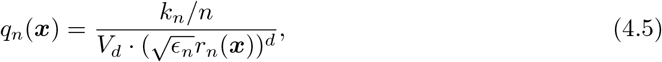

where *V*_*d*_ is the volume of the *d*-dimensional unit ball and *q*_*n*_(***x***) will uniformly converge to the true sampling density *q*(***x***) under mild conditions on *k*_*n*_ and *q* [11]. It is natural to choose

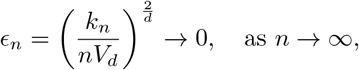

and Theorems A.1 and A.2 in Appendix show that 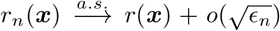 where *r*(***x***) = (*q*(***x***))^−1/*d*^. So we can still denote the kNN kernel as

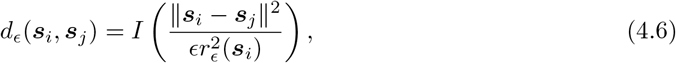

where *ϵ* = *ϵ*_*n*_ → 0 and 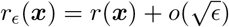.

For the drift part, we will call it the *velocity kernel*. Bearing in mind the intuition that cell *i* is expected to have high probability of transition towards cell *j*, when the corresponding change in gene expression ***δ***_*ij*_ = ***s***_*j*_ − ***s***_*i*_ matches the predicted change according to the velocity vector ***v***_*i*_, we consider the following three schemes of velocity kernels:

- Cosine scheme

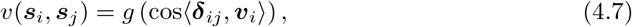
- Correlation scheme

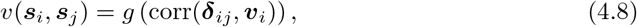
- Inner-Product scheme

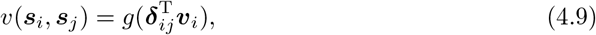

where ⟨***x, y***⟩ and corr(***x, y***) are the angle and the Pearson coefficient between the vectors ***x*** and ***y*** respectively, and *g*(*x*) is a bounded, positive, and non-decreasing function. The overall transition kernel is then defined by

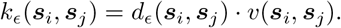

In what follows, we will analyze the continuum limit of the transition kernel *k*_*ϵ*_ for Gaussian diffusion kernel combined with different velocity kernels in Sections 4.2.1–4.2.3, and the analysis for kNN kernel in Section 4.2.4.

#### Remark 4.1.

*We remark that the diffusion kernel does not necessarily bring the diffusion in the final continuum limit of the RNA velocity models as we will see. The use of the name “diffusion” here only respects the convention that it will introduce diffusion type limit in the convergence analysis of graph Laplacians [50].*

#### 4.2.1 Continuum Limit of Cosine Scheme

The transition probability matrix 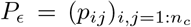 among cells through the Gaussian-cosine scheme is defined by

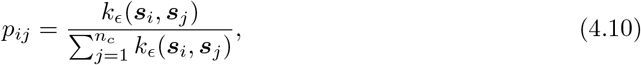

which was utilized in [3].

To study the continuum limit of *P*_*ϵ*_ when the number of samples goes to infinity, we first study the operator 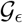 defined by

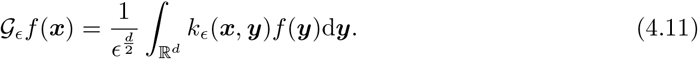

We have the following lemma.

##### Lemma 4.1.

*The operator* 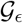 *for Gaussian-cosine scheme has the expansion*

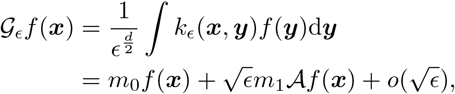

*where*

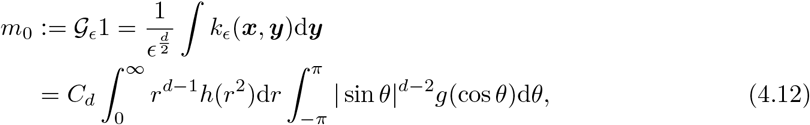

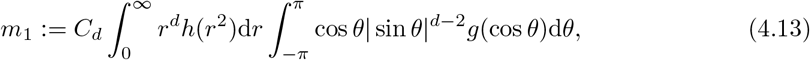

*and*

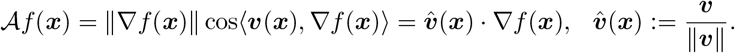

*Here, d >* 1, 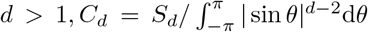 *and S*_*d*_ *is the surface area of the d-dimensional unit sphere.*

*Proof.* Let us assume ***v***(***x***) = ||***v***|| (1, 0, ..., 0)^T^ without loss of generality. The derived result can be transformed back to the original variables by substituting (1, 0, ..., 0) with the vector ***v***(***x***)*/ ||**v**||*.

For simplicity, we first consider the case *d* = 2. Define the polar coordinates

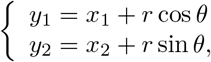

where *θ* is the angle between ***y*** − ***x*** and ***v***(***x***). Then we obtain

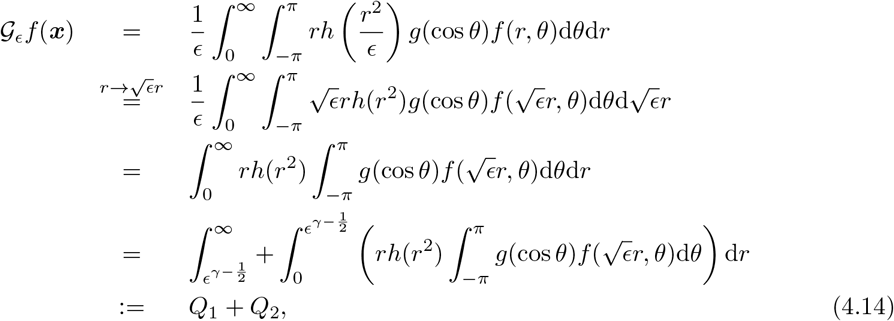

where 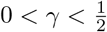. We have

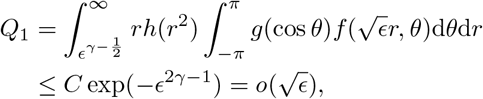

where *C* depends on ||*f*||_∞_ and ||*g*||_∞_ and we utilized the exponential decay of *h*(·) and the inequality 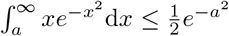.

For the integral in *Q*_2_, by Taylor expansion

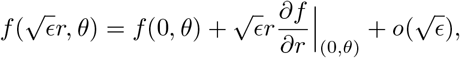

we get

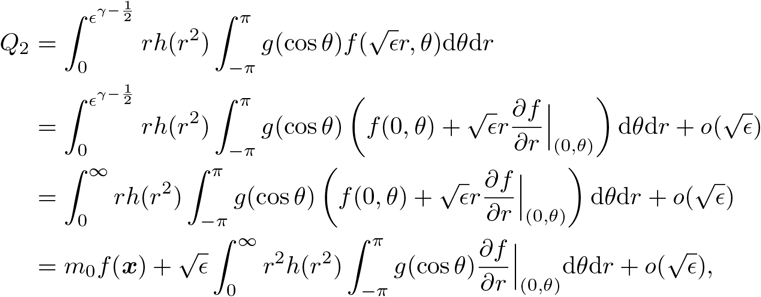

where the extension from the integral domain 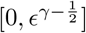 to [0, ∞) will only introduce an exponentially small term by similar argument in estimating *Q*_1_.

For the 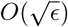 term, note the relation

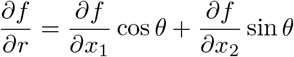

between polar and Euclidean coordinates, we get

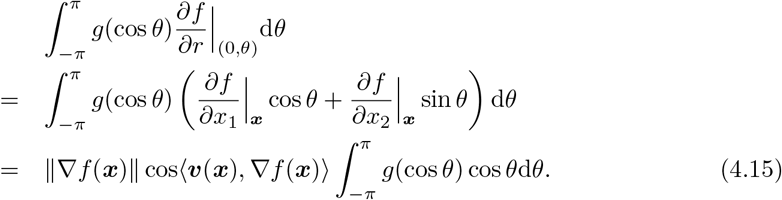

This finishes the proof of the case *d* = 2.

In high dimensions, the derivation is similar. We may consider the coordinate transformation from (*y*_1_, *y*_2_, ..., *y*_*d*_) to (*r, θ, z*_2_, ..., *z*_*d*−2_) defined by

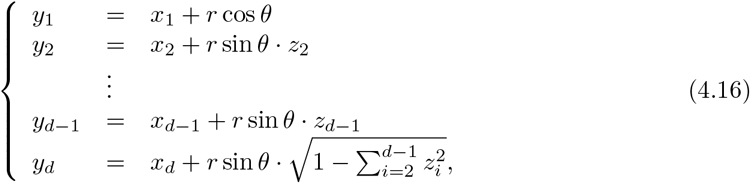

where *r* ≥ 0, −*π* ≤ *θ* ≤ *π*, 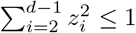. Denote

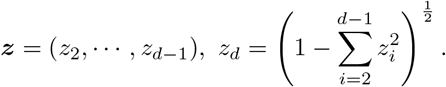

Then the Jacobian

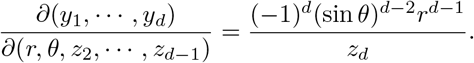

Therefore, we obtain

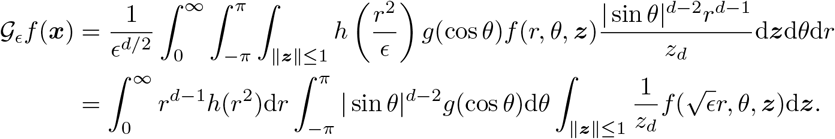

With Taylor expansion

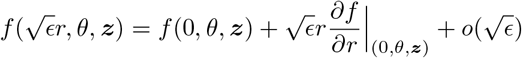

and similar techniques in estimating (4.14) by Laplace asymptotics, we get

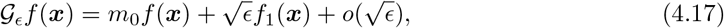

where

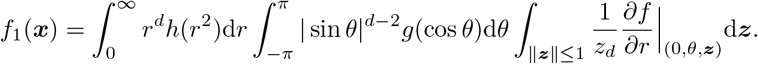

From the relation

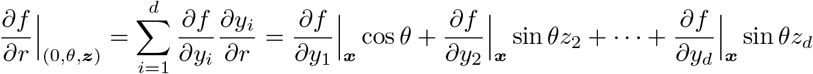

and noting the integral of odd functions with respect to *θ* vanishes, we can simplify the 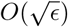 term as

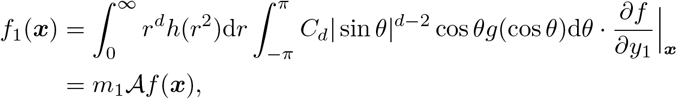

where we used the result 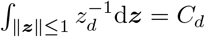.

Given the sample probability density *q*(***x***), then the weighted graph Laplacian has the form

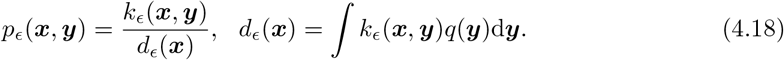

Define the operator

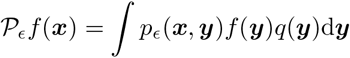

and the generator

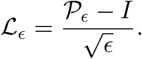

##### Theorem 4.1

(Continuum Limit of the Gaussian-Cosine Scheme). *For the Gaussian-Cosine scheme, we have the limit infinitesimal generator*

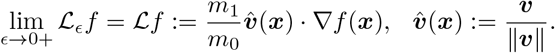

*Proof.* According to Lemma 4.1, we have

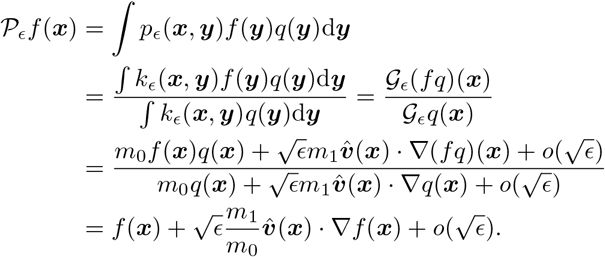

Then the theorem follows obviously.

##### Remark 4.2.

*Theorem 4.1 implies that in the continuum limit, if m*_1_ ≠ 0 *(it holds for general g*(·)*), then the Markov semigroup defined by P*_*ϵ*_ *corresponds to the ODE dynamics*

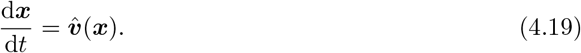

*In this setup, the source and sink states defined by tracing the integral curves of* (4.19) *will be the same as those obtained by solving*

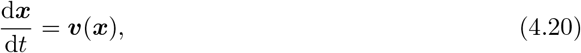

*since the change of the magnitude of the velocity in* (4.20) *does not affect the shape of the integral curves but a reparameterization. However, the transition rule defined by* (4.10) *is not effective on analyzing the landscape and transition behavior among the pluripotent and differentiated cells, which will be addressed in the inner-product scheme.*

##### Remark 4.3.

*The implementation in [3] actually centers both* **δ**_*ij*_ *and v*_*i*_. *So the cosine kernel considered in their paper is equivalent to the correlation kernel below but not the cosine kernel considered here.*

#### 4.2.2 Continuum Limit of Correlation Scheme

The correlation scheme has been utilized in [27, 39]. We have similar expansion to the operator 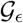 for Gaussian-Correlation scheme.

##### Lemma 4.2.

*The operator* 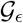 *for Gaussian-Correlation scheme has the expansion*

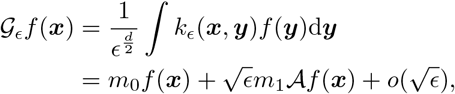

*where*

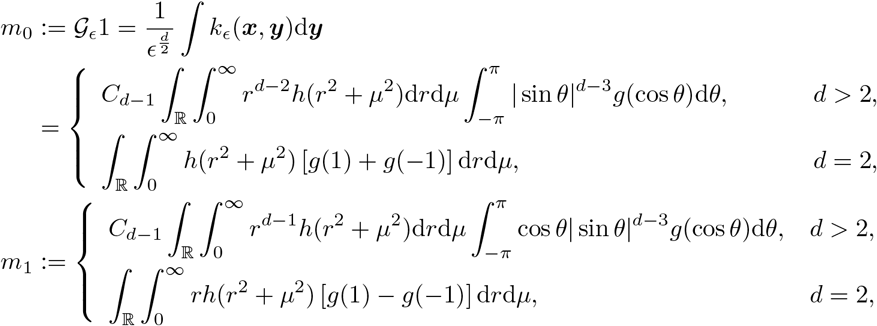

*and*

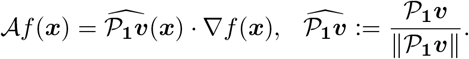

*Proof.* Define projection operator

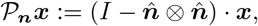

where ***n*** is the normal vector and 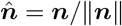. Then the correlation between vectors ***x*** and ***y*** is

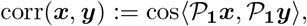

where **1**= (1, ... 1)^T^. It is not difficult to find that the correlation between ***x*** and ***y*** is invariant with respect to the rotations in the hyperplane which is normal to **1**. With this observation, we utilize a new coordinate system which maps

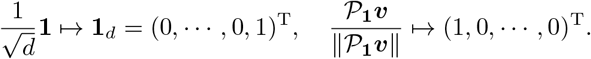

The derived result can be transformed back to the original variable like Lemma 4.1 in a straight-forward way.

In this new coordinate system, the analysis of correlation scheme is similar to the cosine scheme for its first *d* − 1 components. Then we may consider the coordinate transformation below from (*y*_1_, *y*_2_, ..., *y*_*d*_) to (*r, μ, θ, z*_2_, ..., *z*_*d*−2_) when *d >* 2 (the case *d* = 2 is easier and not necessary to do the coordinate transformation since the correlation will be ±1 in this case):

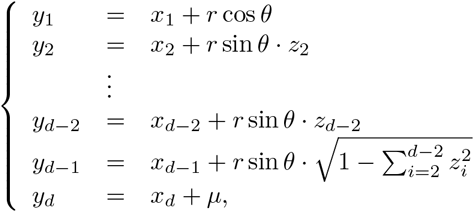

where *r* ≥ 0, *μ* ∈ ℝ, −*π* ≤ *θ* ≤ *π*, 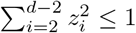. Denote

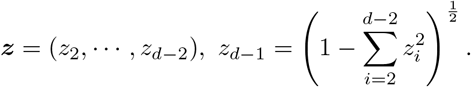

Then the Jacobian

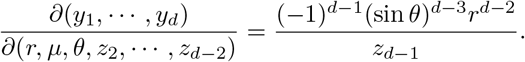

Therefore, we obtain

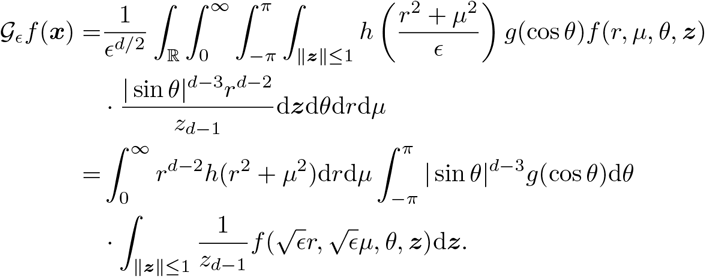

With Taylor expansion

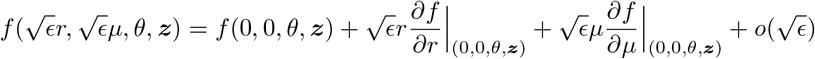

and similar techniques in estimating (4.14) by Laplace asymptotics, we get

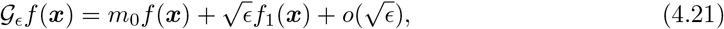

where

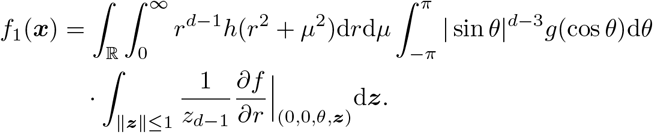

Here we have ignored the odd function with respect to *μ*, then according to the chain rule

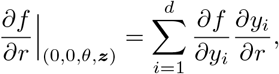

and noting the integral of odd functions with respect to *θ* vanishes, we can simplify the 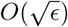 term as

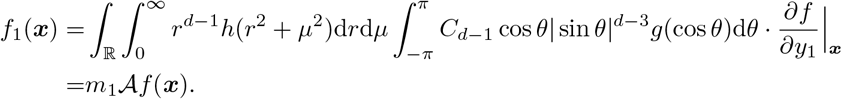

The proof is done.

Then, similar to Theorem 4.1, we get the generator 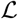.

##### Theorem 4.2

(Continuum Limit of the Gaussian-Correlation Scheme). *For the Gaussian-correlation scheme, we have the limit infinitesimal generator*

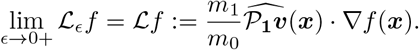

##### Remark 4.4.

*Theorem 4.2 implies that in the continuum limit of correlation scheme, if m*_1_ ≠ 0, *the Markov semigroup defined by P*_*ϵ*_ *corresponds to the ODE dynamics*

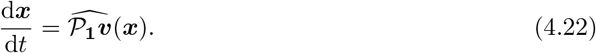

*This means the the effective velocity of the correlation scheme is*

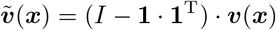

*which will introduce bias into the final result in the identification of root and ending cells. We speculate that the correlation scheme has the possibility to give undesired result in the sense that it is not exactly respect to the inferred velocity.*

#### 4.2.3 Continuum Limit of Inner-Product Scheme

The inner-product scheme is constructed similar to diffusion map [8]. Related idea and methodology has been utilized to analyze the scRNA-seq data analysis [44, 55, 66].

Given the sample probability density *q*(***x***), we define a new kernel

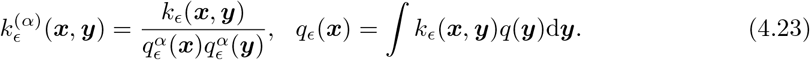

Then we apply the weighted graph Laplacian normalization to this kernel by

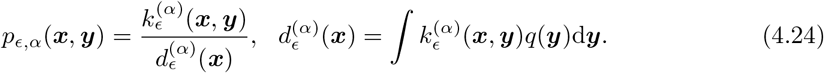

The application of the above construction to the RNA velocity data is straightforward by replacing the density *q*(*x*) with the empirical data distribution of the cell states, i.e.

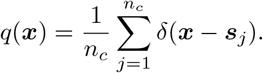

To study the continuum limit in space and time, we define the operator

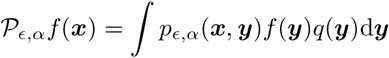

and the generator

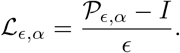

First let us define 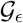 as in (4.11). We have the following lemma.

##### Lemma 4.3.

*The operator* 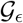 *for Gaussian-Inner-product scheme has the expansion*

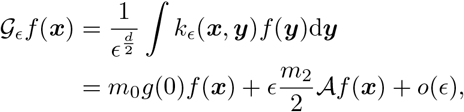

*where the moments*

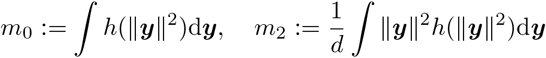

*and*

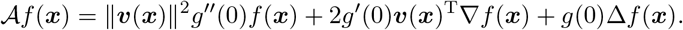

*Proof.* The key idea is to utilize the Laplace asymptotics similar as in estimating (4.14). With inner-product scheme, we have

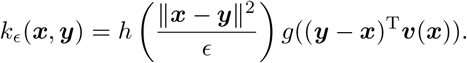

Make the decomposition

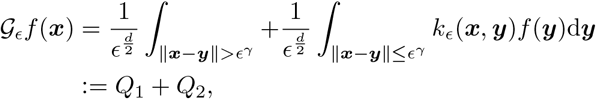

where 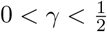. Let 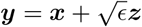. The term

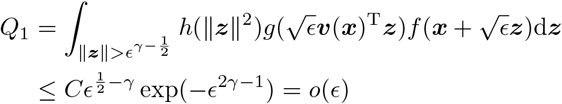

where *C* depends on ||*f*||_∞_ and ||*g*||_∞_, and we utilized the exponential decay of *h*(·) and the inequality 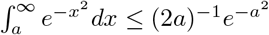 for *a*>0.

For the integrand in *Q*_2_, we have

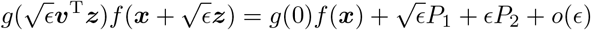

by Taylor expansion, where

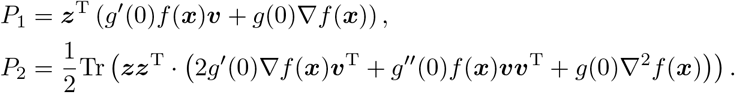

Here Tr(·) is the matrix trace operation.

By the rotation symmetry of ||***z***||^2^ and the fact that the integral of an odd function is zero, we obtain

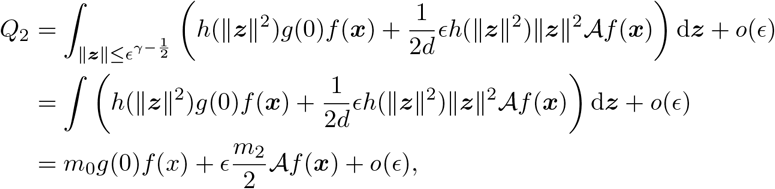

where the extension of the integral domain from 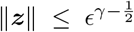 to the whole space will only introduce an *o*(*ϵ*) term by similar argument in estimating *Q*_1_.

##### Theorem 4.3

(Continuum Limit of the Gaussian-Inner-product Scheme). *For the Gaussian-inner-product scheme, we have the limit infinitesimal generator*

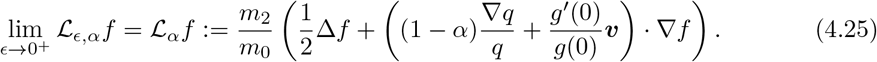

*Proof.* First note that *p*_*ϵ,α*_ in (4.24) is invariant under any multiplicative scaling applied to 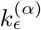. So we can assume in what follows that the normalization *ϵ*^*d/*2^ is implicitly contained in *k*_*ϵ*_(***x, y***), which will not affect the final result.

According to Lemma 4.3, we have the asymptotics

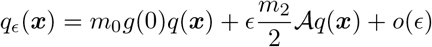

for *q*_*ϵ*_ defined in (4.23). Correspondingly,

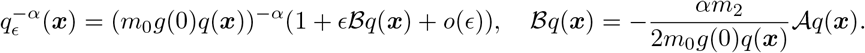

Define

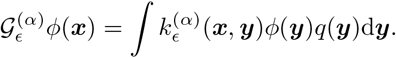

Then

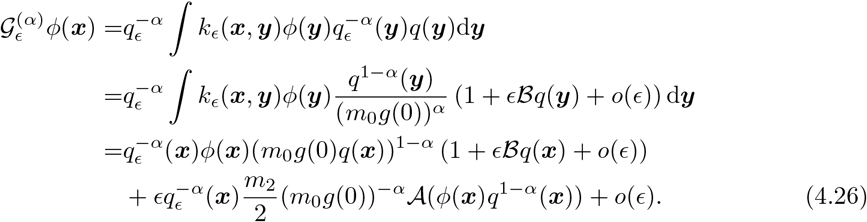

So we obtain

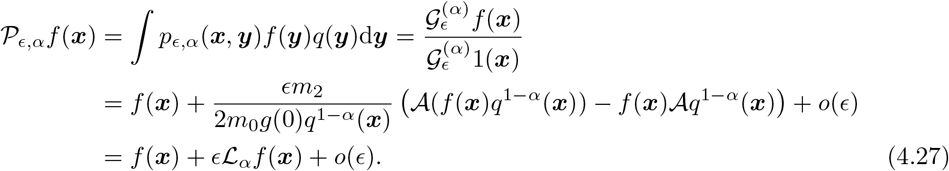

Finally we get

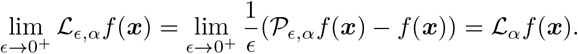

The proof is done.

##### Corollary 4.1.

*If we choose h*(·), *g*(·) *such that m*_2_ = 2, *m*_0_ = 1 *and* 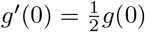, *then*

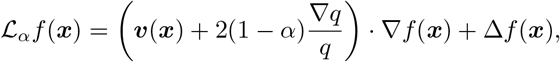

*which is the generator of the stochastic differential equations (SDEs)*

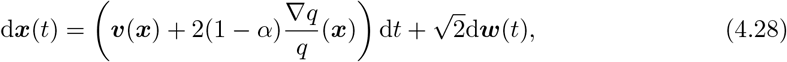

*where **w***(*t*) *is the standard Brownian motion with mean 0 and covariance function* 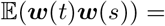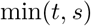. *In general case, similar SDE holds with suitable constants m*_0_, *m*_2_, *g*(0) *and g′* (0).

*Define the data potential*

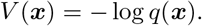

*Then the SDEs* (4.28) *becomes*

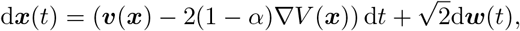

*where the drift term is composed of the gradient part ▽V*(***x***) *from the scRNA-seq sampling distribution, and the non-gradient part from the RNA velocity **v***(***x***). *Specifically, if we choose α* = 1, *then the transition probability will not depend on the data potential V*(***x***) *in the infinite samples limit. This structure opens the way of studying the non-equilibrium steady state and landscape theory for cell developments with scRNA-seq experimental data [65, 63, 17, 44, 55, 66].*

##### Remark 4.5.

*In [4], the authors proposed a class of local kernels to approximate the SDE* (4.2), *which is consistent with our analysis of the inner-product scheme. Note that one typical choice of local kernel is* 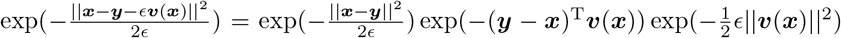, *whose difference with k*_*ϵ*_(*x, y*) *in inner-product scheme is up to O*(*ϵ*) *in g*(·), *incurring no difference in the limit of infinitesimal generator.*

##### Remark 4.6.

*In [27], the transition probabilities among cells were constructed through averaging a RNA velocity-based dynamics and a diffusion-based dynamics. For instance, we consider the transition matrix* 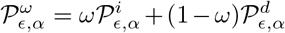 *where* 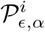 *is the transition probability matrix from inner-product kernel,* 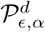 *from pure diffusion kernel and* 0 *< ω <* 1 *is the weight of averaging. Next we provide an understanding of* 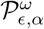 *based on the decomposition of SDE* (4.2) *into the equilibrium and non-equilibrium parts [1, 54, 59, 65].*

*For simplicity, let us assume that the ground-truth dynamics underlying cell-fate development is* 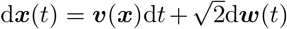 *and the distribution of all data points approximates its steady-state distribution p*^*ss*^(***x***), *which can be guaranteed by the ergodicity of dynamics. With the defined potential landscape ϕ* = −ln *p*^*ss*^, *the velocity term can be decomposed [54] as* ***v***(***x***) = −▽*ϕ*(***x***) + ℓ(***x***), ℓ(***x***) = *J*^*ss*^(***x***)/*p*^*ss*^, *where J*^*ss*^ = ***v***^*pss*^ − ▽*p*^*ss*^(***x***) *is the steady-state probability flux satisfying* ▽ · *J*^*ss*^ = 0*. In terms of the statistical physics interpretation [17], the gradient term ▽ϕ can be viewed as the equilibrium part of velocity, and the curl-like term* ℓ(*x*) *is the non-equilibrium part*.

*With the assumptions above, we apply our results in Corollary 4.1 to the infinitesimal generator of averaging dynamics defined by* 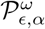, *and conclude that its continuum limit has the form*

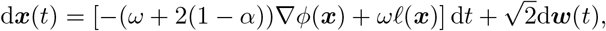

*and the relative proportion of non-equilibrium part ω/*(*ω* + 2(1 − *α*)) *is an increasing function of ω if* 0 *< α <* 1*. Therefore the introduction of weight ω can be understood as tuning the relative weight of equilibrium and non-equilibrium parts of the RNA velocity.*

#### 4.2.4 Implications of kNN Kernels

Sometimes people prefer kNN diffusion kernel with less computational effort due to the sparsity of transition matrix. For example, [3] used kNN-cosine kernel and [39] used kNN-correlation kernel. We will show that for kNN diffusion kernel, the conclusions above still hold with slight difference.

##### Lemma 4.4.

*Suppose q*(***x***) *>* 0 *everywhere. Let r*(***x***) = (*q*(***x***))^−1*/d*^ *be the location-dependent bandwidth function. The operator* 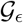 *for kNN-cosine scheme has the expansion*

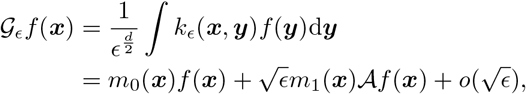

*where*

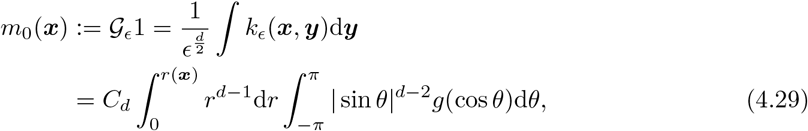

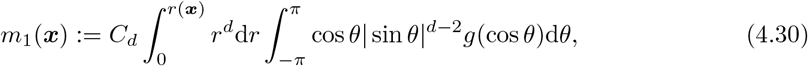

*and*

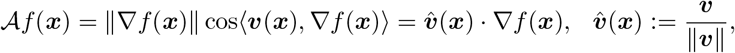

*i.e. we just replace constants m*_0_, *m*_1_ *in Lemma 4.1 with functions m*_0_(***x***), *m*_1_(***x***).

*Proof.* The proof is similar to Lemma 4.1. We only use the case *d* = 2 as an example. Now we have

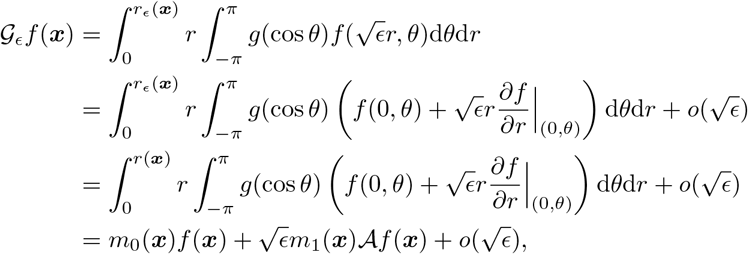

where we utilized the result 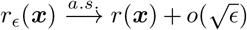 based on the Theorem A.2 in the Appendix.

Now we can derive the limit infinitesimal generator.

##### Theorem 4.4

(Continuum Limit of the kNN-Cosine Scheme). *For the kNN-cosine scheme, we have the limit infinitesimal generator*

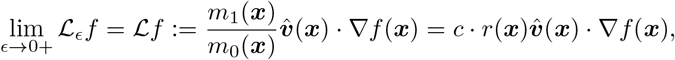

*where c is a constant only related to g*(·) *and dimension d.*

The results for kNN-correlation scheme is similar.

##### Remark 4.7.

*Based upon similar argument in Remark 4.2, the kNN-cosine scheme will give the same root and ending cells produced by Eq.* (4.20) *in the large n limit. The case with q*(***x***) = 0 *is beyond our analysis framework. The kNN-inner-product scheme needs more delicate order analysis on the convergence of r*_*ϵ*_(***x***) *to r*(***x***), *which is considered in the Appendix.*

##### Remark 4.8.

*In [27], the authors combined Gaussian diffusion kernel with kNN kernel, i.e.*

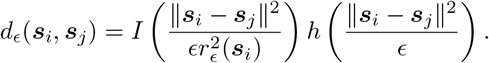

*The overall analysis is similar. We only need to replace m*_0_(***x***) *in* (4.29) *with*

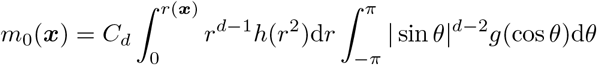

*by inserting h*(*r*^2^) *in the integral of r. The change of m*_1_(***x***) *is similar. The final limit infinitesimal generator is still*

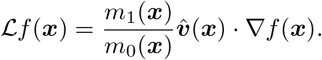

### 4.3 Finding Root and Ending Cells

The root-and-ending cells finding algorithm has been proposed in [3, 27]. The aim of this section is to study its rationale through the continuum limit perspective.

From the derived continuum limits in Theorems 4.1 and 4.3, it is natural to identify the ending cells, i.e. the final differentiated cells, by selecting states with nonzero probability (or higher than a threshold in practice), from the invariant distribution *π* of the transition probability matrix 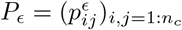:

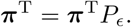

In the limit case as studied in Theorem 4.1, these ending cells corresponding to the absorbing states of the ODE flow map

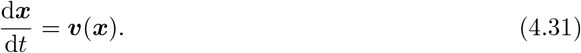

In this case, the limit ODE flow map is not irreducible, thus the invariant distributions are not unique in general. We should start from different, or random initial distributions at the beginning.

The identification of root cells, usually the stem cells, is more subtle. One first defines the backward transition probability matrix 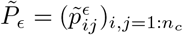 by

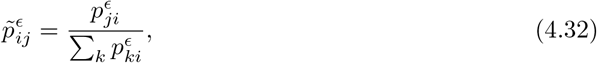

which is the row-normalization of 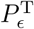; then identify the root cells by selecting states with probability above a threshold in the invariant distribution 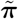 of 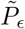.

To motivate the intuition of the above proposal, let us first study the continuous time limit of 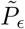 in the discrete states setup.

#### Theorem 4.5.

*If A is the generator of a finite state Markov chain with transition probability matrix P*_*ϵ*_. *Then, the generator of the backward transition probability matrix* 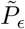 is

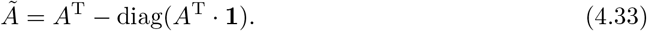

*Proof.* By definition, we have

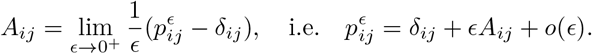

So we get

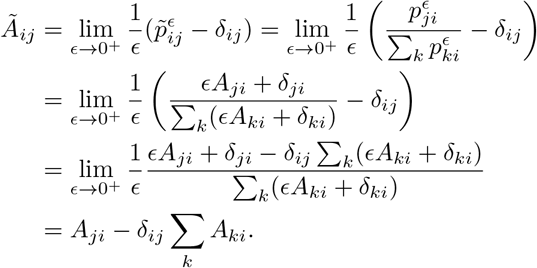

The theorem follows obviously.

Next, we consider the similar limit in continuous states case.

#### Theorem 4.6

(Continuum Limit of the Backward Process). *For transition kernel p*_*ϵ*_(***x, y***) *and operator* 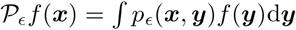, *define the corresponding backward counterparts*

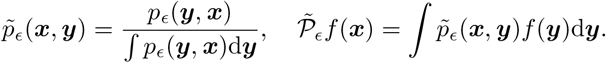

*Then if* 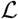 *is the limit infinitesimal generator of* 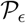 *which satisfies*

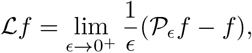

*we have the infinitesimal generator of* 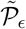

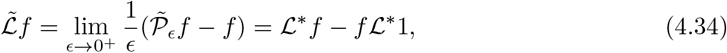

*where* 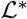 *is the conjugate operator of* 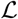, *i.e.* 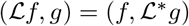.

*Proof.* Consider the operator

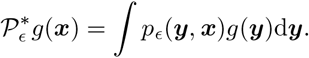

It is straighforward that 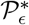 is the conjugate of 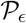, i.e. 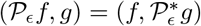, and we have

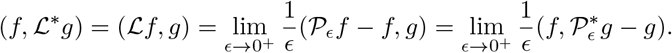

This implies

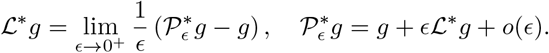

Besides, it is obvious that

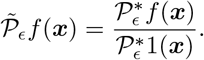

So we have

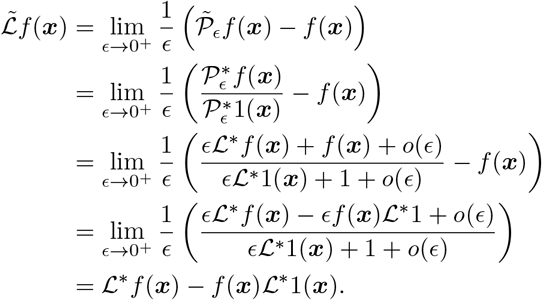

The proof is done.

Now we apply Theorem 4.6 to two most relevant cases.

- ODE case: 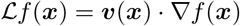. In this case, we have

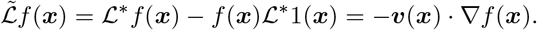 This corresponds to the ODE

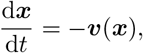

which is exactly the reversed time dynamics of (4.31).
- SDE case: 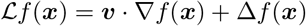. In this case, we have

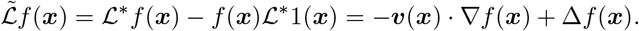 This corresponds to the SDEs

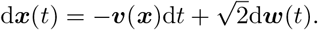 To apply the theorem to the scRNA-seq data with transition rules (4.10) and (4.24), we need to take into account the data distribution *q*(*x*). Define

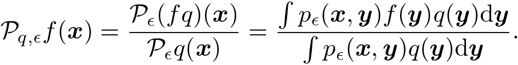 It is not difficult to show that

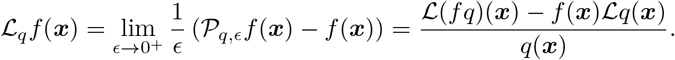 Correspondingly, for

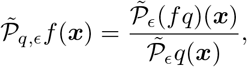

we have

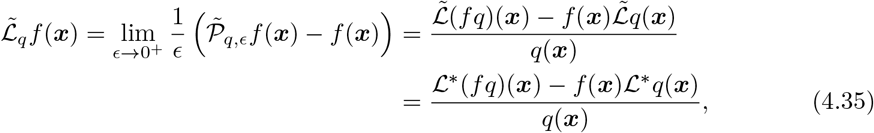

where we utilized the conclusion in Theorem 4.6. From Theorem 4.1 we know that for the cosine scheme with Gaussian diffusion kernel, 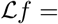 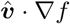 for appropriate *h*(·) and *g*(·). Then its conjugate operator 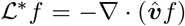. Therefore, we have

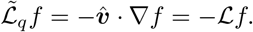 Similarly, for correlation scheme, from Theorem 4.2 we also have

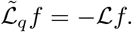 For the inner-product scheme, from Lemma 4.3 we know that

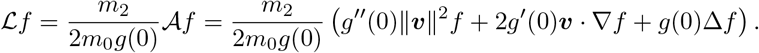 Then its conjugate operator is

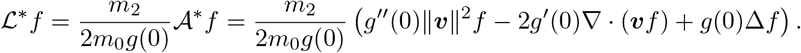 Similar to Theorem 4.3, simply replacing 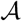 with its conjugate 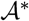 in the proof, we can show that

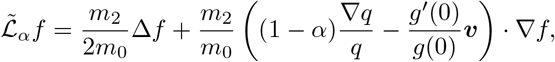

which is similar as the operator 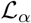 (4.25) except reversing the direction of velocity ***v***.

Overall, the above derivations show that for all of cosine, correlation and inner-product schemes, if we replace the transition kernel with its backward form, their continuum limit will follow the ODE or SDE dynamics, by reversing the direction of the RNA velocity ***v***. Similar results also hold for kNN-cosine or kNN-correlation (or Guassian-kNN-cosine/correlation) schemes with similar derivations. This gives the rationale of the identification of root and ending cells through backward and forward transition rules, respectively.

### 4.4 Finding Development Routes: Transition Path Theory

With the root and ending cells detected by RNA velocity, the next question is to ask how the cell state evolves along the development trajectories that connect the starting and target cell fates. Unlike the conventional picture of trajectory inference (such as pseudotimes), the dynamics revealed by RNA velocity might be more complex, because of local fluctuations, rotations and oscillations, as well as multiple sources and sinks along the trajectory. Except for calculating the most probable transition paths in the continuous set-up [39], the majority of existing tools opt to visualize the trajectories with local velocity arrows or connecting streamlines in the reduced-dimensional space, where a more quantified and global description of development path is needed.

We reason that the transition path theory [15] might be a good candidate, which has been established for general Markov process such as diffusion [14], jump [33] and Markov chains [34], and yielded fruitful applications in molecular dynamics and chemical reactions [5, 32]. Our proposed method to find development routes can be understood as a discrete, data-driven version of the continuous approach described by [39]. Below we only focus on the theoretical aspect of our proposal, whose algorithmic details will be discussed in the continued work [28]. We will mainly follow [32] for the illustration of the transition path theory, and ignore most proofs of the theorems which can be referred to [32] for details.

#### 4.4.1 Coarse-graining of Transition Dynamics

The rapid growth of scRNA-seq data size poses computational challenges to the downstream analysis. Therefore we propose an optional step here to first coarse-grain the transition dynamics on the scale of clusters instead of single-cells to reduce the computational complexity.

##### Definition 4.1

(Coarse-graining of Markov Chain). *Given an ergodic, microscopic Markov chain* {*x*_*t*_} *on the state space S with transition probability matrix P* = {*p*(*x, y*)} *and a partition 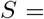 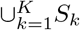 the coarse-graining of* {*x*_*t*_} *is defined as a Markov chain* {*X*_*t*_} *on the state space* 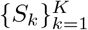 *with transition probability matrix*

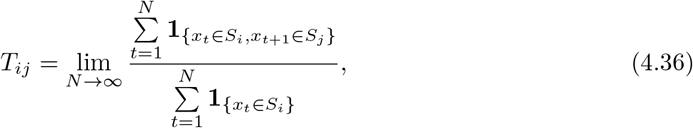

*where the limit is taken in the almost sure sense, and* **1**_{·}_ *is the indicator function with* **1**_{*exp*}_ = 1 *if the logical variable exp* =*TRUE, and 0 otherwise.*

We remark that the naive upscaling of the microscopic Markov chain {*x*_*t*_} by considering the induced transition *X*_*t*_ on the coarse-grained space at each step is not valid since *X*_*t*_ defined in this way is not necessarily Markovian. This is related to the well-known lumpability concept in Markov chain theory [13, 25]. Here we take alternative viewpoint by defining the coarse-grained chain through the time average limit instead of the single step transition.

The partition of the state space can be achieved by the clustering of cells, or by the simultaneous reduction of multi-scale dynamics [13, 66]. The coarse-grained transition probability matrix can be indeed calculated analytically instead of through numerical simulations:

##### Proposition 4.1.

*The coarse-grained transition probability matrix defined by* (4.36) *can be expressed as*

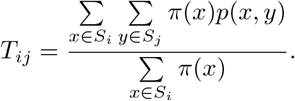

*Proof.* Consider the stochastic process *y*_*t*_ = (*x*_*t*_, *x*_*t*+1_), which is indeed a Markov chain with stationary distribution 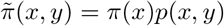. Then from the ergodic theorem for *y*_*t*_ we have

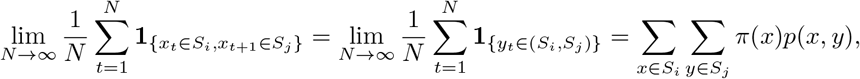

and for *x*_*t*_ we have 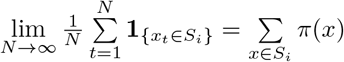.

It is also straightforward to verify tha t the coarse-grained Markov chain *X*_*t*_ with {*T*_*ij*_} has the stationary distribution ***μ*** with 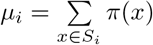.

For simplicity of notations to present the transition path, below we take the index of {*X*_*t*_} as integer set ℤ, where *t* = 0 corresponds to the interested time point, and the minus time points represent the trajectories prior to *X*_0_. With this setup, *X* is stationary and *X*_*t*_ obeys the invariant distribution always.

#### 4.4.2 Defining Transition Paths and Their Probabilities/Fluxes

From the forward and backward transition approach described in Section 4.3, we are able to identify the sets of root and ending clusters, and denote them as committor starting set *A* and target set *B*, respectively. Note that both *A* and *B* may contain several states, corresponding to the complex dynamics of multiple root or ending states and various connecting trajectories. One advantage of transition path theory indeed lies in the quantification of such dynamics.

To begin with the derivation of transition path theory, we first define the core concepts of in-transition times and transition paths as follows.

##### Definition 4.2

(In-Transition Times). *For a given path* {*X*_*t*_}, *the in-transition times from set A to B are defined as the union of sets*

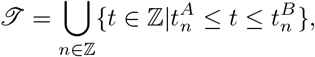

*where 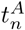 and 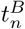 are the nth exit and entrance time of set A and B respectively such that*

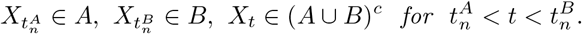

##### Definition 4.3

(Transition Paths). *For a given path {X*_*t*_}, *the nth transition path from A to B is*

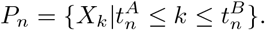

The set of all transition paths is defined as 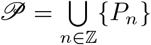.

We are interested in quantifying the probability distribution of transition paths ensemble, which is defined as:

##### Definition 4.4

(Probability of Transition Paths). *The probability of observing transition path at state i is defined as*

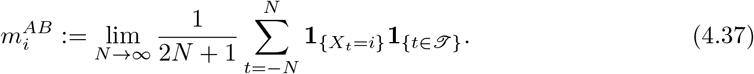

Intuitively, 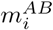 describes the likelihood that the cell is on a transition path from *A* to *B* and bypassing state *S*_*i*_. To compute 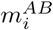, we need the notion of committor functions and their associated first entrance/last exit time.

##### Definition 4.5

(First Entrance and Last Exit Time). *Given the path {X*_*t*_} *of stationary process X, the first entrance time* 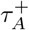 *into set A, and the last exit time* 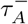 *from set A are defined as*

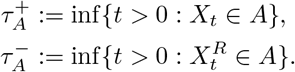

*where 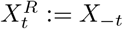 is the path of time-reversed process of X*_*t*_, *and* 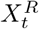 *has the transition probability matrix* 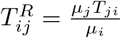.

##### Definition 4.6

(Committor Function). *The forward and backward committor functions are defined as*

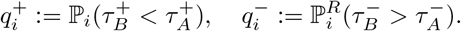

*Here* ℙ_*i*_ *denotes the probability of the forward process X conditioned on X*_0_ = *i and* 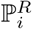 *the probability of the reversed process X*^*R*^ *conditioned on* 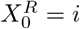.

From the definition, we can also interpret 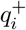 as the probability that the cell starting from cluster *S*_*i*_ first enters set *B* rather than set *A*, and 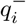 the probability that the cell arriving at cluster *S*_*i*_ came last from set *A* instead of *B*. The committor functions provide a natural soft clustering of the states by measuring the affinities with starting set *A* or target set *B*.

To compute the committor functions, we can derive the linear equations they satisfy.

##### Proposition 4.2.

*The committor functions solve the following discrete Dirichlet problems*

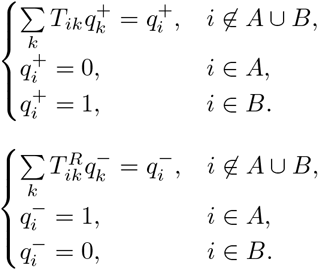

*Proof.* Define the Markov chain 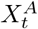 with absorbing set *A* through the transition probability

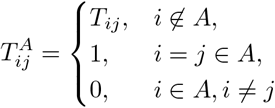

and the hitting time 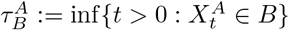 of 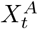 into set *B*. Then we have 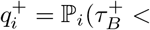 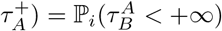. From the Markov property of 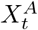, for *i* ∈ (*A ∪ B*)^*c*^ we have

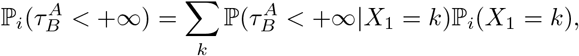

which yields the equation for 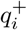, since 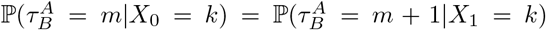 for *k* ∈ (*A ∪ B*)^*c*^. The equation for 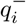 is similar. The validation of remaining boundary conditions are straightforward from the definition of committor function.

With committor functions, we have the following representation of transition paths probability.

##### Proposition 4.3.

*The probability of transition paths defined in* (4.37) *can be expressed as*

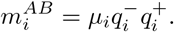

The intuition of the above expression is clear. To observe the transition paths at state *i*, we pick it with the stationary distribution *μ*_*i*_, and require that the path last exit from set *A* and first enter the set *B*. This happens with the probability 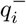 and 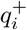, respectively.

##### Remark 4.9.

*The proportion ρ*^*AB*^ *of the time that a cell spends on the transition paths from A to B has the form*

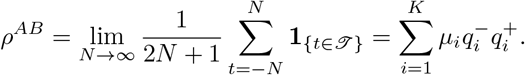

Similarly, we can define the probability flux of transition paths, which is important to the detection of development routes discussed below.

##### Definition 4.7

(Probability Flux of Transition Paths).

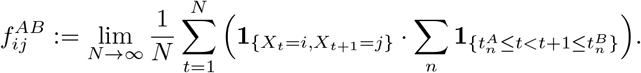

Roughly it tells the proportion of cells that are on a transition path from *A* to *B* and moving directly from *S*_*i*_ to *S*_*j*_. We can also write *f*_*ij*_ in terms of committor functions, which also serves as the numerical strategy for computation.

##### Proposition 4.4.

*The probability flux of transition paths can be expressed as*

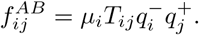

#### 4.4.3 Finding Development Routes via Transition Paths Flux

Trajectory inference aims to indicate how the state of cells evolve in a step-wise way. We therefore define the concept of development routes to illustrate this physical picture.

##### Definition 4.8

(Development Route). *A development route ω*_*dr*_ = (*i*_0_, *i*_1_, *..., i*_*n*_) *from set A to B is a path connecting A and B without self-interactions (loops) such that*

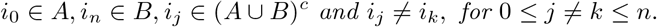

To quantify development routes of Markov chain *X*_*t*_, we need to eliminate the effect of detours along transition paths, and therefore define the notion of effective current (or net flux) based on the probability flux of transition paths.

##### Definition 4.9.

*The effective current of transition paths from state i to j during the transition from set A to B is defined as*

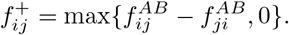

With the effective current, we can specify the capacity of each development route.

##### Definition 4.10

(Capacity and Bottleneck). *Given the development route ω*_*dr*_ = (*i*0, *i*1, ..., *i*_*n*_) *from set A to B, its capacity is defined as*

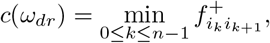

*and the bottleneck is defined as the corresponding edge with minimal effective current.*

The underlying intuition of the definition can be indeed understood by an analogy with the stream in water pipes or traffic on the freeways. The capacity of water pipes or freeways, is limited by the narrowest point where the minimal amount of water stream or traffic can pass through. Similarly, the transition of cell state from *A* to *B* is constrained by the bottleneck on the development route.

For all the possible development routes, we can calculate their capacity and consider routes with larger capacity, which corresponds to the more dominant trajectories during the transition. The ranking of all development routes capacity can be done effectively using an iterative edge-removing strategy [33]. We leave the details of algorithmic implementation of transition path theory to find development routes based on RNA velocity in our continued work [58].

## 5 Conclusion

The introduction of RNA velocity allows the prediction of future states in single-cell RNA sequence (scRNA-seq) data and yields fruitful results to reveal the dynamics of actual development process [46], while several theoretical issues regarding the models and analysis of RNA velocity remain to be elucidated. In this paper, we have proposed a mathematical framework to investigate the modeling, inference and downstream analysis aspects of RNA velocity.

Here we presented both the deterministic and stochastic models of RNA velocity, and derived the analytical solutions for both models. Particularly, we provided the expression for the exact probability distribution at any time in stochastic model. With the introduced models and analytical solutions, we then revisited the algorithms to infer parameters in RNA velocity model, and proposed an EM algorithm for the newly-derived complete stochastic model through maximum likelihood estimation.

Next, we dedicated to the theoretical issues on downstream analysis, particularly focusing on recovering the dynamical system models from RNA velocity. We derived the continuum limits of various constructed cell-cell transition dynamics from RNA velocity, which depended on the choice of velocity kernels (cosine, correlation or inner-product) and diffusion kernels (Gaussian or kNN). Our analysis revealed that while the cosine scheme in velocity kernel uncovers the streamlines in the deterministic ODE dynamics of RNA velocity model, the inner-product scheme corresponds to the stochastic dynamics described by SDE, which can incorporate the transitions among meta-stable states. Meanwhile the correlation scheme is associated with the deterministic dynamics with altered velocity that might be potentially unwanted. Through the delicate analysis on kNN kernel, we also proved that the choice of kNN over Gaussian in the diffusion kernel did not affect the overall continuum limit except for the prefactors.

Based on our analysis, we then validated the rationale to find root and ending cells of previously proposed “forward and backward diffusion” strategy [27], from the continuous dynamical system aspects. It is shown that the difference between forward and backward transitions in the continuum limit only lies in the reversal of velocity direction. Finally, we proposed a method to infer the development routes from RNA velocity-based transition rules, which was specialized to cope with the complex dynamics of multiple root/ending states and various connecting path. We demonstrated the derivation of the method from transition path theory for Markov process.

Compared with previous stochastic models of RNA velocity that focused on moment equations of the chemical master equation [3, 39], the analysis of our proposed stochastic model featured for the derivation of an exact solution for the probability evolution. Such probability expression is especially useful to derive most-likelihood estimators of parameters for the stochastic model.

Currently, there exist two major streams of research to infer and analyze the underlying dynamics of scRNA-seq data. The first class of *data-driven* approaches, which are well represented by pseudo-time inference methods [20, 38, 51] for snapshot data, seek to construct development trajectories from the intrinsic manifold representation of data. On the other hand, the second class of *model-based* approaches as exemplified by SCUBA [31], Pseudo-dynamics [16] and Waddington-OT [41], aim to connect the probability distribution of single-cell data with a continuous dynamical system model, and are widely applied in the analysis of time-series sequencing data. Weinreb et al. [55] have shown theoretically that the fundamental limitations of snapshot data make the data-driven methods incapable of revealing the complex, non-equilibrium dynamics accurately. Meanwhile the model-based proposals may encounter difficulties in solving the high-dimensional Fokker-Planck equations numerically and infer the large amount of parameters in the model. Here we provide the mathematical justifications that the discrete cell-cell transition dynamics constructed from RNA velocity, even for snapshot data, can indeed converge into the continuous dynamical system model (not necessarily equilibrium) via the large sample limit, and we also propose the method to dissect the complex trajectories of such dynamics with solid theoretical guarantee in the well-established transition path theory.

In the future, we anticipate that the RNA velocity model can be further improved by taking the genetic interactions into account [12], and customizing for the time-series scRNA-seq data. Overall, our mathematical analysis of RNA velocity in this paper provides the starting point to develop further models and methods in a more rational and consistent way.

## Acknowledgment

T. Li and Y. Wu is supported by the NSFC under grants Nos. 11421101 and 11825102, and Beijing Academy of Artificial Intelligence (BAAI).

## Appendix. The kNN Radius Estimate

In this Appendix, we will prove a useful kNN radius estimate as *n* → ∞, which is summarized in Theorems A.1 and A.2. Although some classical results have been obtained in the literature concerning the consistency of the kNN density estimate [11, 29], the asymptotic expansion of the kNN radius is seldom discussed. Our analysis is more direct and independent of the result considered in [50]. It is not sharp, but enough for our continuum limit analysis of RNA velocity kernels. We leave further delicate order estimate as a future work.

Consider a random vector ***X*** with smooth density function *q*(***x***). Given *n* data points, we can estimate the density function at *x* through kNN approach

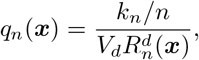

where *V*_*d*_ is the volume of the *d*-dimensional unit sphere, *R*_*n*_(***x***) is the kNN radius defined as the distance to the *k*_*n*_th nearest neighbor of ***x***. Without loss of generality, we set ***x*** = **0** and denote *R*_*n*_ = *R*_*n*_(**0**). By definition, *R*_*n*_ obeys the order statistics with density

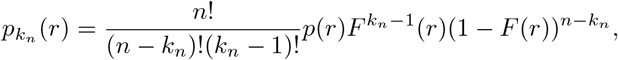

where *p*(*r*) is the density of the radius *R* = ||***X***||:

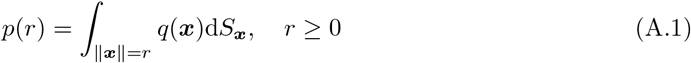

and *F* is its distribution function 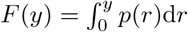.

Below we will first estimate the expectation of *R*_*n*_ to achieve an intuition about its scale. We assume *q*(**0**) *>* 0 all through the analysis.

### Lemma A.1.

*Under the condition k*_*n*_ → ∞, *k*_*n*_/*n* → 0 *and the assumption*

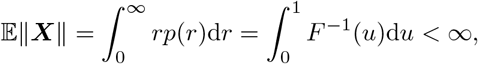

*we have*

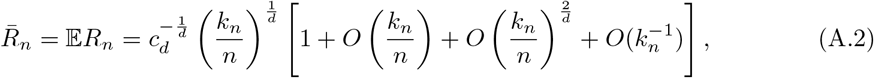

*where c*_*d*_ = *V*_*dq*_(**0**).

*Proof.* For any 0 *< ϵ* ≤ 1, define

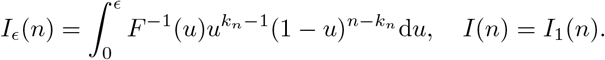

Then we have

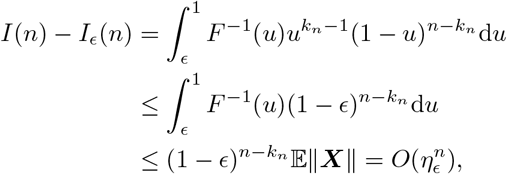

where *η*_*ϵ*_ is a generic constant belongs to (0, 1), which can be taken as 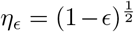 in the current step as long as *k*_*n*_ *n/*2.

Now let us consider *I*_*ϵ*_(*n*). We need to estimate the order of *F*^−1^(*u*) when *u* is small. For small *δ >* 0, we have

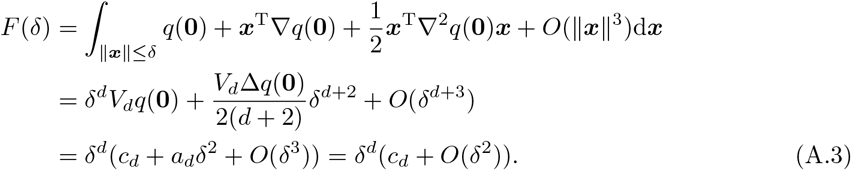

Denote *G*(*u*) = *F*^−1^(*u*), then *G*(*δ*^*d*^(*c*_*d*_ +*O*(*δ*^2^))) = *δ.* Let *u* = *δ*^*d*^(*c*_*d*_ +*O*(*δ*^2^)). We get *δ* = *O*(*u*^1*/d*^), and

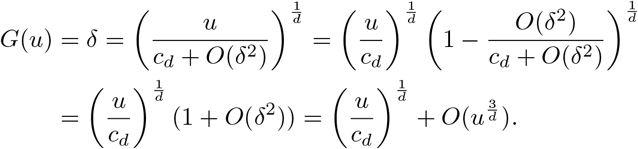

Therefore we obtain

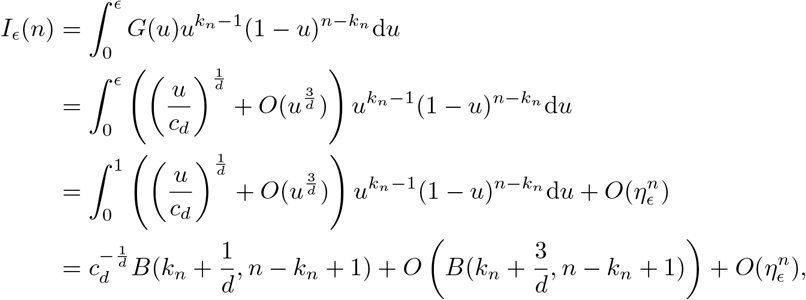

where 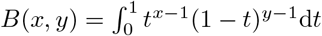 is the beta function. Utilizing the asymptotics

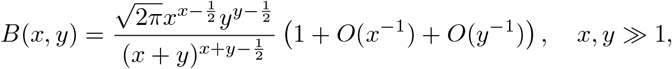

we obtain

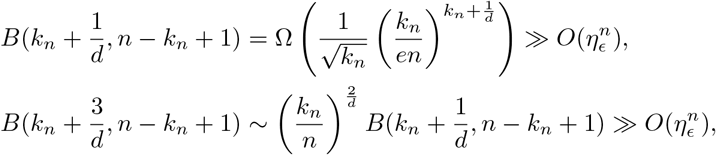

where Ω(·) is the asymptotic lower bound function and we have used the condition *k*_*n*_*/n* → 0. Therefore, we get

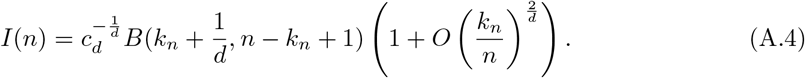

Then the expected radius

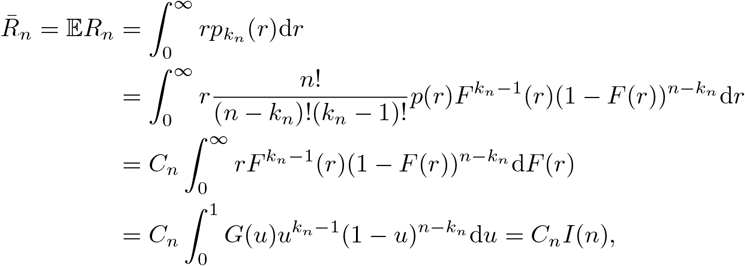

where *C*_*n*_ = *n*!/(*n*−*k*_*n*_)!(*k*_*n*_−1)!. Using Stirling’s formula and (A.4) we obtain (A.2) immediately.

Lemma A.1 suggests that the kNN radius *R*_*n*_(***x***) has the form *R*_*n*_(***x***) = *h*_*n*_*r*_*n*_(***x***) with the scale

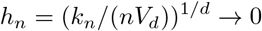

and the location dependent bandwidth

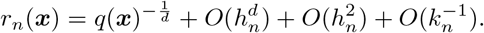

Below we first establish the convergence in leading order in almost sure sense.

### Theorem A.1.

*If k*_*n*_/*n* → 0 *and k*_*n*_/ ln *n* → ∞, *then*

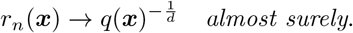

*Proof.* Set ***x*** = **0** without loss of generality. According to the Borel-Cantelli lemma, we only need to prove that for any small *ϵ >* 0,

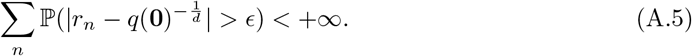

Denote 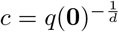. We have

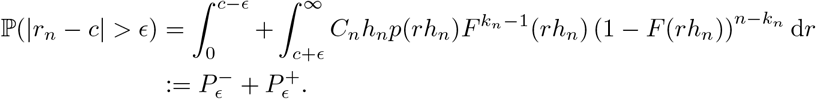

From (A.3), we have the asymptotics

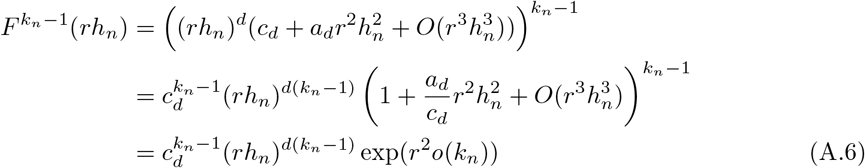

and

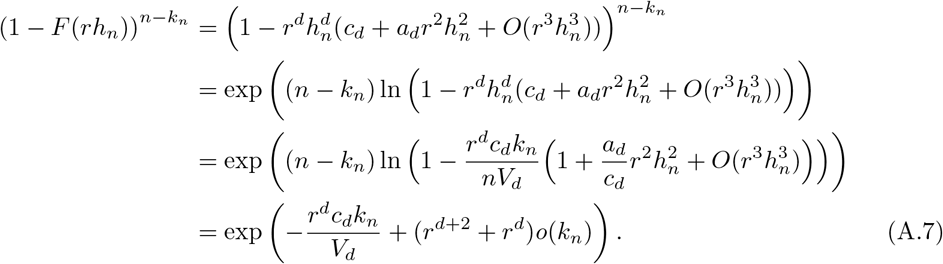

Define 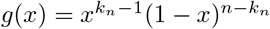. Then (*k*_*n*_ − 1)/(*n* − 1) is the unique maxima of *g*, and *g* is increasing when 0 *< x <* (*k*_*n*_ − 1)/(*n* − 1) and decreasing when (*k*_*n*_ − 1)/(*n* − 1) *< x <* 1. Utilize (A.3), we can easily show that

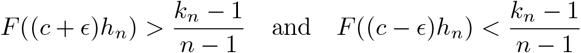

for small *ϵ* and big enough *n*. Then using (A.6) and (A.7) and Stirling’s formula, we obtain

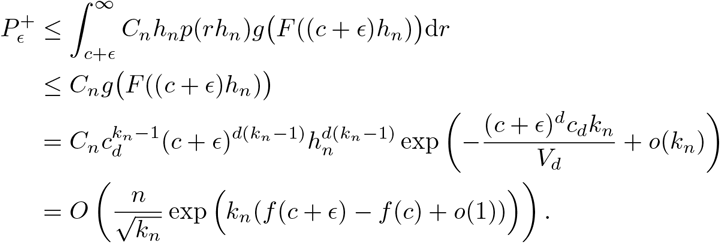

where

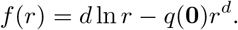

The function *f* is increasing when *r < c* and decreasing when *r > c*.

With similar estimate for 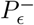, we obtain

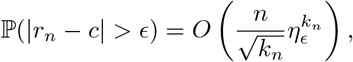

which is *o*(*n*^−2^) under the condition *k*_*n*_/ ln *n* → ∞. Then (A.5) follows and the proof is done.

### Theorem A.2.

*If k*_*n*_/*n* → 0 *and k*_*n*_/*n*^*α*^ → ∞ *for some α >* 2/(*d* + 2), *then for d >* 1, *we have*

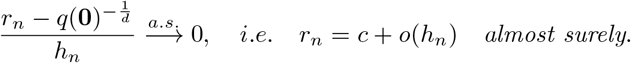

*Proof.* Similar to Theorem A.1, now we consider

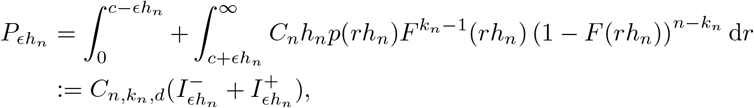

where 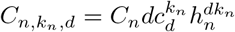 which has the asymptotics

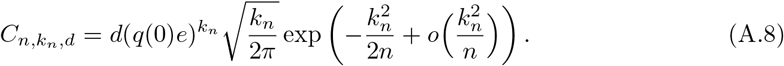

According to the proof of Theorem A.1, for any *δ >* 0 and *n* big enough, we have

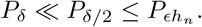

Therefore, to estimate 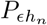, we only need to estimate 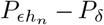. In what follows we will mainly use the Laplace’s method to estimate 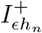 since the estimation of 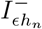 is similar. First we have

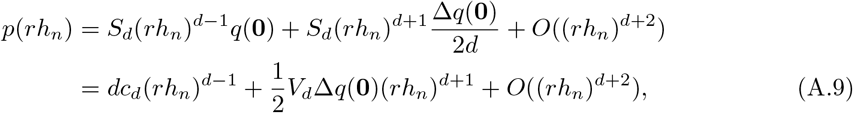

where *S*_*d*_ is the surface area of the *d*-dimensional unit sphere. Utilizing (A.6), (A.7) and (A.9), we get

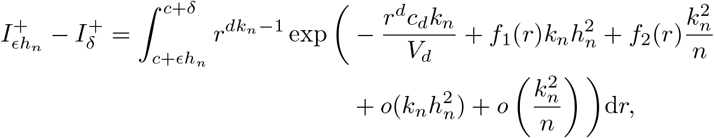

where

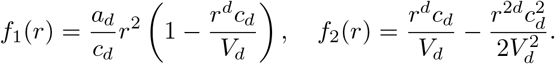

We can verify that *f*_1_(*c*) = 0 and *f*_2_(*c*) = 1/2. By the arbitrary smallness of *δ*, we have

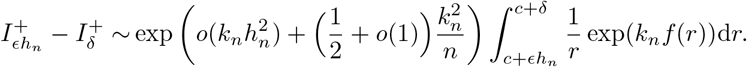

Denote

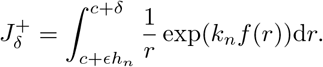

For *c* + *ϵh*_*n*_ < *r < c* + *δ*, there exists *ξ >* 0 such that

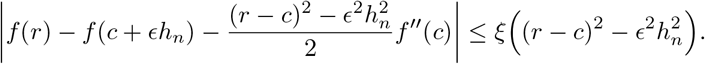

We have

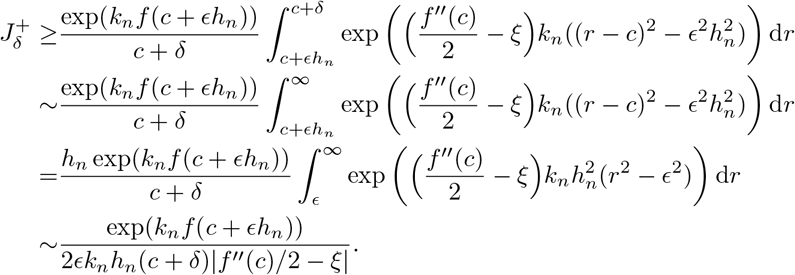

Similar upper bound can be obtained, so we obtain

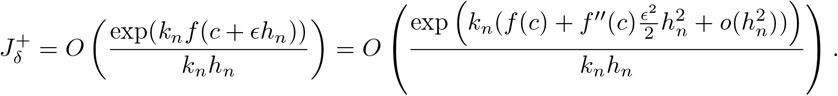

Therefore,

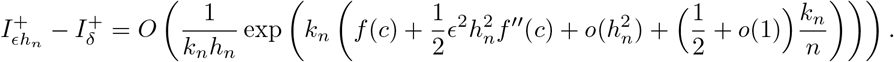

With similar estimate for 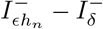, we get

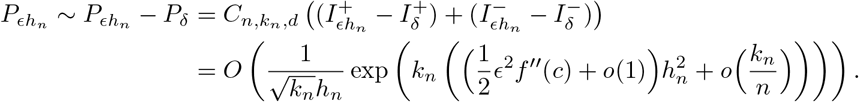

If *d >* 1, then 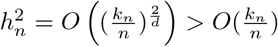. Therefore

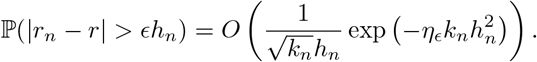

Under the condition that *k*_*n*_/*n*^*α*^ → ∞ for some *α >* 2/(*d* + 2), we have 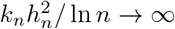, so we get the desired estimate.

## Notes

### Competing Interest Statement

The authors have declared no competing interest.

